# Temporal and Spatial Scales of Human Resting-state Cortical Activity Across the Lifespan

**DOI:** 10.1101/2025.03.28.645952

**Authors:** John Bero, Colin Humphries, Yang Li, Aviral Kumar, Heungyeol Lee, Maxwell Shinn, John D. Murray, Timothy J. Vickery, Daeyeol Lee

**Author notes:** **Address for Correspondence:** Daeyeol Lee, Timothy J. Vickery.

## Abstract

Sensorimotor and cognitive abilities undergo substantial changes throughout the human lifespan, but the corresponding changes in the functional properties of cortical networks remain poorly understood. This can be studied using temporal and spatial scales of functional magnetic resonance imaging (fMRI) signals, which provide a robust description of the topological structure and temporal dynamics of neural activity. For example, timescales of resting-state fMRI signals parsimoniously predict a significant amount of the individual variability in functional connectivity networks identified in adult human brains. In the present study, we quantified and compared temporal and spatial scales in resting-state fMRI data collected from 2,352 subjects between the ages of 5 and 100 in Developmental, Young Adult, and Aging datasets from the Human Connectome Project. For most cortical regions, we found that both temporal and spatial scales decreased with age throughout the lifespan, with the visual cortex and the limbic network consistently showing the largest and smallest scales, respectively. For some prefrontal regions, however, these two scales displayed non-monotonic trajectories and peaked around the same time during adolescence and decreased throughout the rest of the lifespan. We also found that cortical myelination increased monotonically throughout the lifespan, and its rate of change was significantly correlated with the changes in both temporal and spatial scales across different cortical regions in adulthood. These findings suggest that temporal and spatial scales in fMRI signals, as well as cortical myelination, are closely coordinated during both development and aging.

**Significance Statement:** Temporal and spatial scales of resting-state cortical activity in humans measured by fMRI largely decreased throughout the lifespan, except that for some regions in the prefrontal cortex they peaked similarly during adolescence. In addition, whereas cortical myelination consistently increased throughout the lifespan, its variation across different cortical networks and the rate of age-related changes were correlated with the dynamics of temporal and spatial scales of rs-fMRI activity, suggesting that the spatio-temporal scales of cortical activity and cortical myelination might be co-regulated during development and aging.

## Introduction

The human brain goes through massive changes throughout the lifespan that manifest as development and decline in cognitive functions. Sensorimotor systems tend to mature first in early childhood followed by language and other high-order cognitive skills (Casey et al., 2005; Gogtay et al., 2004; Sowell et al., 2004). Fluid abilities, such as working memory, show a steady decline from early adulthood, whereas crystalized abilities, such as semantic memory, improve and then plateau around age 60 (Murman, 2015; Harada et al., 2013; Salthouse, 2010; Levy, 1994). These different phases of cognitive development and aging are a result of changes in the underlying anatomical and cytoarchitectural properties of the brain. Brain size, cortical thickness, and myelination increase during the early stages of life, followed by the loss of cortical tissues (Stiles and Jernigan, 2010; Bethlehem, et al., 2022; Blinkouskaya, et al., 2021). However, these anatomical changes are not uniform, exhibiting a regional heterogeneity in different development phases, which could be the main driver for the differences in cognitive time courses throughout the lifespan (Casey et al., 2005; Stiles and Jernigan, 2010). Similarly, functional connectivity (FC) computed using resting-state fMRI (rs-fMRI) also exhibits age-related changes. For example, FC during childhood is dominated by short-range local interactions, which are gradually replaced by long-distance interactions in adulthood (Fair et al., 2009). Aging studies have demonstrated the opposite effect, with networks exhibiting decreased connectivity as the brain matures (Geerlings et al., 2015). This trajectory implies a peak in development sometime between childhood and young adulthood (Betzel et al., 2014; Coupé et al., 2017; Bethlehem, et al., 2022; Edde et al., 2021).

Despite these previous findings, however, many aspects of cortical functions during development and aging remain poorly understood. For example, elementary statistical properties of the rs-fMRI signal, such as temporal and spatial autocorrelations, vary within and across individuals (Murphy et al., 2013; Friston et al., 1995; Burt et al., 2020; Shinn et al., 2023). Temporal and spatial scales in the fMRI signals might reflect important biological factors, including molecular (Burt et al., 2020; Huang et al., 2018), structural (Sethi et al., 2017; Fallon et al., 2020), activity state (Fascianelli et al., 2019; Arbabshirani et al., 2019) and network (Honey et al., 2012; Song et al., 2014) properties of the brain. Therefore, identifying the factors that influence the intrinsic temporal and spatial timescales of rs-fMRI signals is important to fully understanding the brain’s dynamics throughout lifespan. Nevertheless, due to the restricted age ranges used in previous studies on the topic, the complete age profile of spatial and temporal scales in the human brain has not been fully characterized (Raut et al., 2020; Ito et al., 2020; Shinn et al., 2023; Han et al., 2022; Truzzi and Cusack, 2023; Gao et al., 2020; Wu and Gollo, 2025).

In the present study, we estimated the timescale of the rs-fMRI signals using the correlation coefficient between the signals in two successive time steps. In addition, we estimated the spatial scale of the rs-fMRI signals and its regional variation by measuring how steeply the FC decays with distance over the cortical surface. We examined how these temporal and spatial scales of rs-fMRI signals as well as cortical myelination change from infancy into late adulthood by examining multiple datasets spanning the ages from 5 to 100 years from the Human Connectome Project. We found that the spatial and temporal scales of rs-fMRI have similar non-monotonic changes for some prefrontal regions, peaking during adolescence followed by a decrease for the rest of the lifespan. Furthermore, we found that age-related changes in temporal and spatial scales of rs-fMRI signals were correlated with the level of myelination across different cortical areas during childhood, but with the rate of changes in myelination during adulthood. These results suggest that the coordination between spatio-temporal scales of cortical activity and myelination might have important implications for brain development and aging.

## Materials and Methods

### Datasets and participants

The data analyzed in the present study were taken from three different datasets of the Human Connectome Project (HCP); developmental dataset (HCP-D) composed of youth and adolescents (age 5∼22y), 1,200 subject dataset (HCP-YA) composed of young adults (age 22∼37y), and Aging dataset (HCP-A) composed of middle aged and elderly adults (age 37∼100y). In the following, we will refer to these three datasets as Child, Adult, and Old age groups, respectively. We analyzed the preprocessed resting state fMRI time series in these datasets. Subjects were excluded based on their average framewise displacement (FD), computed as the sum of the absolute values of the differentiated realignment estimates (by backwards differences) at every time point (Power et al., 2012). Based on the distribution of mean framewise displacement, a value corresponding to the distribution mean plus two times the standard deviation was used as a cutoff threshold. The final sample consisted of 627 Child subjects, 1,036 Adult subjects, and 689 Old subjects, for a total of 2,352 subjects (1,151 males, 1,201 females; Table 1). As described below, there were some variations in data collection methods, quality control procedures and preprocessing steps across the datasets. In addition, the sampling density (number of subjects/age) was not uniform across ages and was maximal for the Adult dataset.

**Table 1.** Demographic Information for subjects.

| Dataset | Age Range | Original Sample | FD Threshold (mm) | Final Sample | Males/<br>Females | Age Mean ± SD (years) |
| --- | --- | --- | --- | --- | --- | --- |
| Child | 5 - 22 | 652 | 0.340 | 627 | 290/337 | 14.58 ± 4.04 |
| Adult | 22 - 37 | 1,084 | 0.273 | 1,036 | 558/478 | 28.72 ± 3.68 |
| Old | 37 - 100 | 725 | 0.378 | 689 | 303/386 | 60.05 ± 15.63 |

### Data Acquisition

Child and Old data were collected on a 3-T SIEMENS Prisma scanner, and included T1w anatomical (multi-echo MPRAGE, resolution=0.8 mm × 0.8 mm × 0.8 mm, TR/TI/TE = 2,500/1,000/1.8 ms, flip angle = 8°), T2w anatomical (variable-flip-angle TSE, resolution = 0.8 mm × 0.8 mm × 0.8 mm, TR/TE = 3,200/564 ms, turbo factor = 314) and rs-fMRI scans (GRE EPI, resolution = 2 mm × 2 mm × 2 mm, TR/TE = 800/37 ms, measurements = 488, MB factor = 8, flip angle = 52°, scan time = 6.30 min). Adult data were collected on a custom 3-T SIEMENS Skyra scanner, and included T1w anatomical (MPRAGE, resolution = 0.7 mm × 0.7 mm × 0.7 mm, TR/TI/TE = 2,400/1,000/2.4 ms, flip angle = 8°), T2w anatomical (variable-flip-angle TSE, resolution = 0.7 mm × 0.7 mm × 0.7 mm, TR/TE = 3,200/565 ms, GRAPPA = 2) and rs-fMRI scans (GRE EPI, resolution = 2 mm × 2 mm × 2 mm, TR/TE=720/33 ms, measurements = 1,200, MB factor = 8, flip angle = 52°, scan time = 14.4 minutes).

### Data analysis

#### Preprocessing

Structural and functional MRI data were processed using the HCP minimal preprocessing pipelines as described previously (Glasser et al., 2013; Bero et al., 2023). The T1w and T2w images were aligned with the anterior commissure-posterior commissure (AC-PC) line, brain extracted, corrected for intensity bias and registered to the MNI template. The white matter and pial surfaces were traced, and the data were projected onto the cortical surface via FreeSurfer (Fischl, 2012). Myelin content was estimated by taking the log ratio of T1w/T2w signal intensities on a voxel-by-voxel basis (Glasser and Van Essen, 2011; Ganzetti et al., 2014) on the standard CIFTI grayordinate surface. The resting-state fMRI data underwent motion correction, susceptibility distortion correction, brain extraction, registration to MNI and anatomical space, time series normalization, component-based denoising using ICA-FIX and high pass filtering (0.0005 Hz). Processed volumetric data were projected onto the standard CIFTI grayordinate surface (59,412 voxels) and smoothed with a full-width half max of 2 mm. Other details of the processing were identical to those used in previous studies (Glasser et al., 2013; Glasser et al., 2016; Harms et al., 2018).

#### Postprocessing

To facilitate comparisons between datasets, some common processing steps were applied to all final outputs following their individual preprocessing pipelines. All functional surface maps were parcellated based on the Schaefer parcellation, which is a functional parcellation based on resting-state functional connectivity data with 400 total regions of interest (ROI) separated into 17 resting-state networks (Figure 1a; Schaefer et al., 2018). In addition, a stricter high-pass filter was applied to all time series at 0.008 Hz, and all ROI time series were standardized prior to analysis. Since the current study focuses on regional variation, global patterns and correlations were removed in all our data at the subject level for the main results presented in the present study. The global signal for each frame was computed as the average of signals across all ROI, and global signal regression was performed for each participant (Murphy and Fox, 2017; Li et al., 2019). Nevertheless, we also tested how the estimates of temporal and spatial scales of rs-fMRI signals are affected by comparing the results obtained with and without the global signal regression.

**Figure 1.**
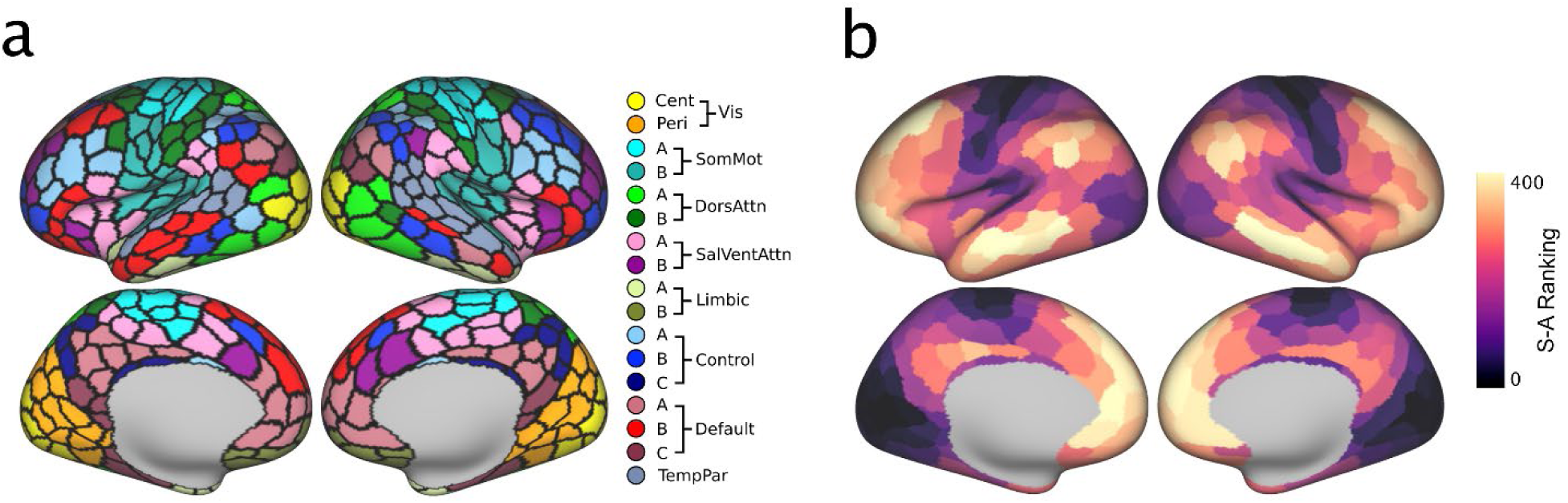
Brain maps for resting-state functional networks and sensorimotor-association axis. (a) Functional networks identified in Schaefer (2018) parcellation. (b) Ranking of ROI in Schaefer parcellation along the sensorimotor-association (S-A) axis identified by Sydnor et al. (2021).

#### Analysis of Temporal and Spatial Scales

To estimate the timescale of rs-fMRI signals, we computed the first-order (lag-1) autocorrelation of each parcellated time series, namely, the Pearson correlation between the blood-oxygen-dependent-level (BOLD) signals in two successive time bins. We also computed the lag-1 autocorrelation in the global signal for each subject. For a first-order autoregressive process, this lag-1 autocorrelation (lag-1 TA, or ϕ_1_) is related to its time constant (τ) as τ = −TR/ln ϕ_1_, and can efficiently characterize the individual variability in the temporal autocorrelations in the rs-fMRI signals (Shinn et al., 2023). Lag-1 autocorrelation is also correlated with multiple temporal features of BOLD signals including the so-called long-memory dynamics (Shafiei et al., 2020; Shinn et al., 2023). To compare these long-memory or long-range processes to lag-1 autocorrelation, we also estimated the fractional integration constant *d* from an ARFIMA model by using a univariate wavelet-based Whittle estimator, which has been used in previous studies to capture long memory dynamics (Achard et al., 2008; Sela and Hurvich, 2009; Achard and Gannaz, 2015).

To determine the spatial scale, a distance metric must first be established on the cortical surface. For this study, geodesic distances were computed on the pial brain surfaces from the centroids of each Schaefer ROI, creating a 400 × 400 distance matrix. Functional connectivity matrices were computed by taking the Pearson correlation between each pair of parcellated time series (also with dimensionality of 400 × 400). For a given ROI, all other brain regions were binned according to their geodesic distance, using 1-mm bins, and the mean correlation with each bin was used to generate a spatial autocorrelation (SA) function for that ROI. Then, the resulting SA was fit by an exponential function with 2 parameters, the rate at which the correlation falls off with distance (SA-λ) and the baseline (SA-∞), using the method used in Shinn et al. (2023), as follows.

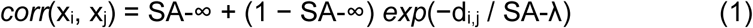

where x_i_ is the time series for region i and d_i,j_ is the distance between regions i and j. An ADAM optimizer (a variation of stochastic gradient descent) parallelized with 9 CPU cores was used to estimate the model parameters. We used SA-λ, akin to a length constant, as a measure of spatial scale in the rs-fMRI signals. Network averages were obtained by averaging the parameters of all ROIs within a network. To test the reliability of the resulting brain map of the temporal and spatial scales, we computed average brain maps in two randomly divided subgroups in each age group and computed the average correlations between them over 100 repetitions.

#### Age Dependence of Spatial and Temporal scales

To quantify the age-related change in spatial and temporal scales of rs-fMRI signals, we first used linear regression to generate brain maps of age coefficients for spatial and temporal scales in each age group. In addition, statistical significance of the age effect was tested using a linear mixed effects model with age and network coded as continuous and discrete variables, respectively. The statistical significance of overall age-network interaction was assessed by comparing a full model (AC = Age + Network + Age × Network) and a reduced model (AC = Age + Network) using a χ^2^-test test (Chen et al., 2013). In the full model, individual age-network interactions were tested using the difference relative to the mean as contrast vectors. For visualization of age-related changes in the spatial and temporal scales over the entire lifespan, curves were fit separately for the network averages in each age group. This was done using a locally weighted scatterplot smoothing (LOWESS) regression model (with a fraction of 0.5), which is not influenced by the uneven sampling distributions as present in the data (Cleveland, 1981). Separate fits were performed for each age group to remove any artificial features that might arise due to differences in acquisition or preprocessing between the datasets. Significance of non-monotonic trends in each age group were tested using a 2-tailed t-test on the second-order age term from a regression model with a quadratic term for each network (p<0.05) with the requirement that the maximum or minimum of the model must fall within the age range of the dataset in question. Both spatial and temporal scales estimated in this study revealed some discontinuities at the age boundaries between different datasets, apparently due to the differences in the scanning protocol and pre-processing methods. We did not apply any methods to harmonize these results, since the age distributions in different datasets did not overlap. This also allowed us to preserve the original physical units for spatial and temporal scales. We used the same methods to analyze age-related changes in the global signal.

### Comparison with other brain maps

We constructed brain maps corresponding to the mean values and age coefficients for temporal and spatial scales of rs-fMRI signals for each ROI in the Schafer parcellation. To examine age-related changes in cortical microstructures, similar maps were also created for cortical myelination using the log ratio of T1w/T2w as a proxy (Glasser and Van Essen, 2011; Ganzetti et al., 2014) and were analyzed using the same mixed effects model and the LOWESS regression fit used to analyze temporal and spatial scales. We compared these brain maps as well as the ranking of each ROI along the sensorimotor-association (S-A) axis that accounted for many anatomical and functional features of the human cortex (Figure 1b; Sydnor et al., 2021, https://pennlinc.github.io/S-A_ArchetypalAxis/). The statistical significance of correlations between different brain maps was determined using a t-test.

## Results

### Age-related changes in timescales of fMRI signals

We first computed the lag-1 autocorrelation to examine how the timescales of rs-fMRI signals in different cortical areas changed across different age groups (Figure 2a). The resulting brain maps of timescales were robust in all age groups, as indicated by the correlation coefficient between the maps generated for randomly divided two halves in each group (r = 0.82, 0.78, and 0.73, for Child, Adult, and Old, respectively). The timescales were relatively short in the insular cortex and cingulate gyrus, and long in the occipital and parietal regions (Figure 2a). These brain maps were highly similar across different age groups at the network level (r > 0.95, p < 0.001; Figure 2b and 2c). We also found that the whole-brain mean timescales significantly decreased from Child to Adult to Old age groups (mean ± SD = 0.59 ± 0.20, 0.41 ± 0.19, and 0.39 ± 0.20, for Child, Adult, and Old, respectively; 1-way ANOVA, p<0.05; Figure 2b, 2c), which presumably resulted from the differences in imaging protocols across datasets. These results were nearly identical when we used the long-memory parameter *d* estimated with the ARFIMA model instead of the lag-1 autocorrelation. Indeed, the long-memory parameter and lag-1 autocorrelation were highly correlated across networks for all age groups (r > 0.99, p < 0.001).

**Figure 2.**
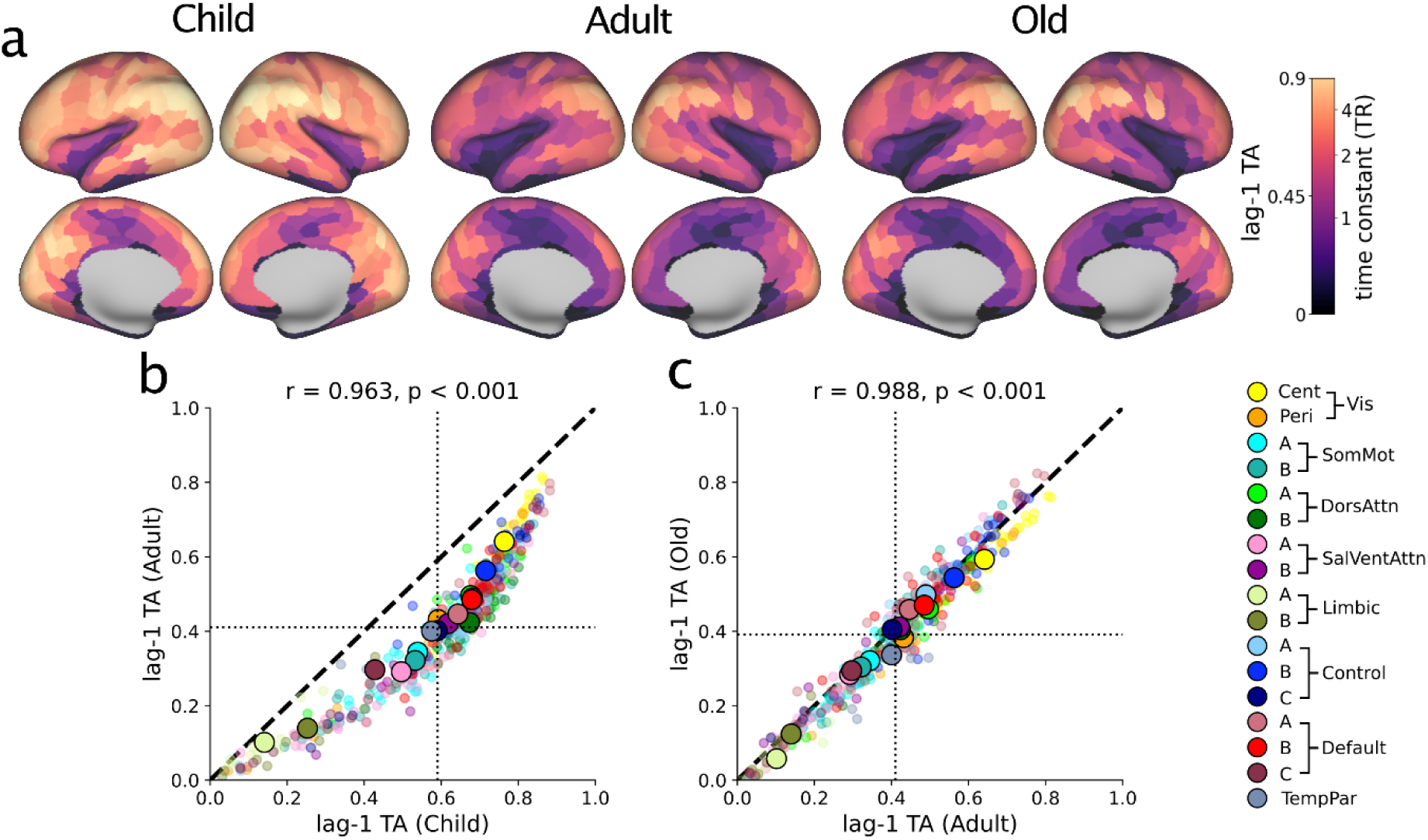
Timescales of rs-fMRI activity. (a) Brain maps of mean timescale (lag-1 TA) maps for Child (left), Adult (center), and Old (right) age groups. (b) and (c) Scatter plots of the timescales for Child vs Adult (b) and Adult vs Old (c) age groups. Small and large symbols show the values for individual ROI and network averages. Dotted lines show group means.

Overall, we found that the timescales tended to decrease for most FC networks in all age groups, as indicated by the significantly negative age coefficient in a linear mixed effects model (Table 2). We also computed the age coefficient of a simple linear regression model for each ROI and found that they are negative in most ROI, except that some ROIs around Brodmann area 10 in the Child age group showed significantly positive age coefficients (p<0.05; Figure 3a). In addition, we found that for all age groups, the full model with age × network interaction terms fit the data better than a reduced model without them, suggesting that the slopes of individual network trajectories were not uniform within each age group. We found that the trajectory of the timescale was more complex and heterogeneous in the Child age group than in the Adult and Old age groups. Most FC networks showed significant age × network interaction in the Child age group (n=11 out of 17), whereas this was the case for fewer networks in Adult (n=4) and Old (n=6) age groups (Table 2). The Limbic-A, Limbic-B, and Default-C networks showed significantly less negative (i.e., more positive) slopes than the average in both Adult and Old age groups.

**Figure 3.**
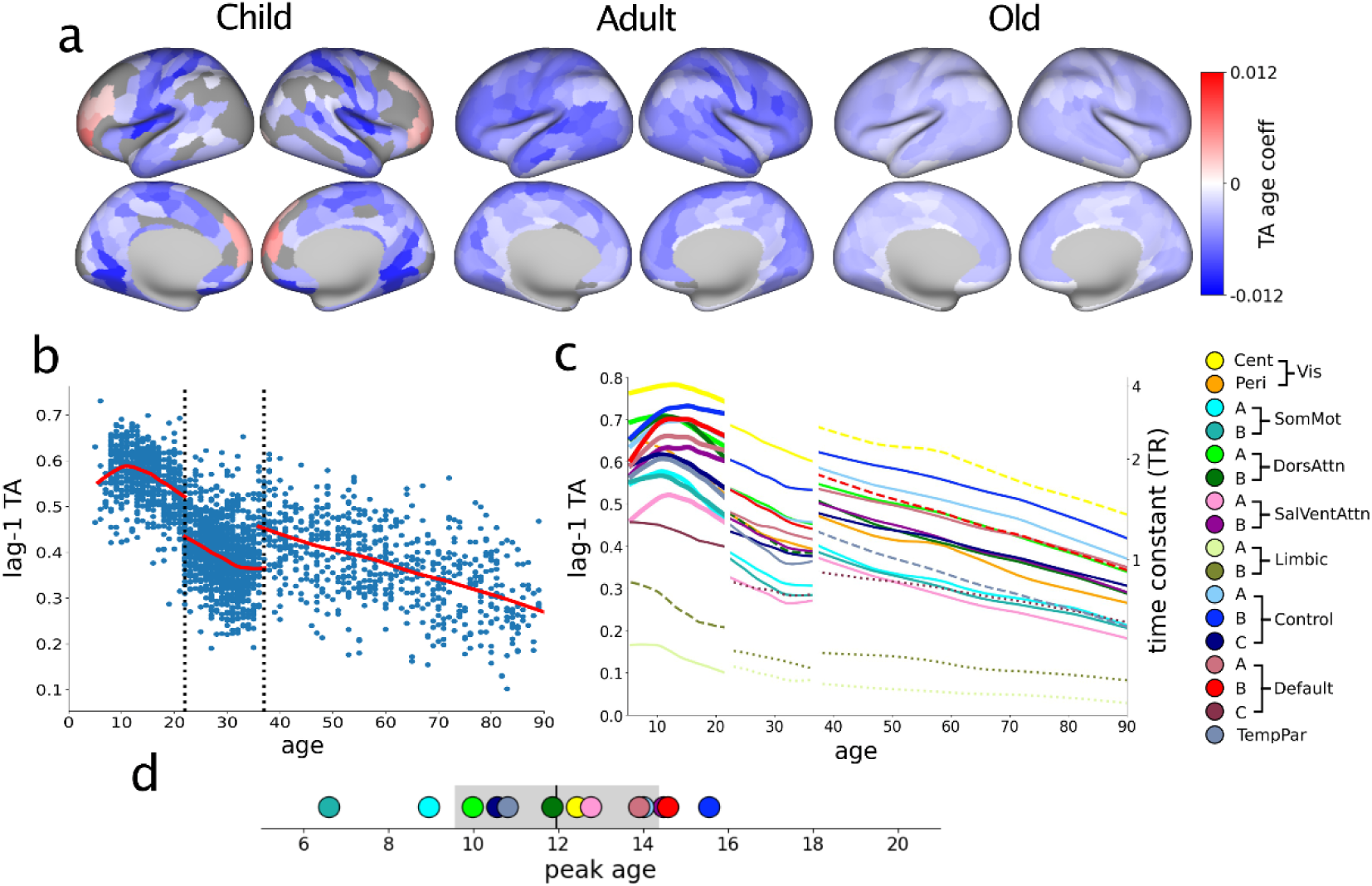
Age-related changes in the timescale of rs-fMRI signals. (a) Brain maps of the age coefficient for timescale in Child (left), Adult (middle), and Old (right) age groups. Gray regions indicate the ROI in which the age effect was not statistically significant (p>0.05). (b) Scatter plots of the average timescale (lag-1 TA) for the entire cortex across individual subjects. The red line represents the LOWESS fit and vertical dotted lines show the border between age groups. (c) LOWESS fit for timescale in each FC network. Dotted/dashed lines show networks that are significantly more positive/negative than the average (see also Table 2). (d) Peak ages for timescales in networks with significant non-monotonic time course in the Child age group. The black vertical line and gray bar correspond to the mean±SD.

**Table 2.** Age-related changes in timescale, spatial scale, and cortical myelination across cortical networks estimated by linear mixed effects models for different age groups. In addition to the F-value of the full model and the age coefficient, the list of networks with significant positive and negative age×network interactions (Bonferroni-corrected for multiple comparisons) and the result from the χ^2^-test for the difference between the full model and the reduced model without interaction terms are shown. *, p<0.05.

| | Group | F | Age | Positive | Negative | $\chi^2$ (p) |
| --- | --- | --- | --- | --- | --- | --- |
| Timescale | Child | 1,548* | -0.0045* | Vis-C<br>SV-B<br>Cont-A, Cont-B<br>Def-A, Def-B<br>(N=6) | Vis-P<br>SM-A, SM-B<br>DA-B<br>Lim-B<br>(N=5) | 308.4*<br>($p < 10^{-10}$) |
| | Adult | 1,321* | -0.0056* | Lim-A, Lim-B<br>Def-C<br>(N=3) | DA-B<br>(N=1) | 75.0*<br>($p < 10^{-8}$) |
| | Old | 778* | -0.0033* | Lim-A, Lim-B<br>Def-C<br>(N=3) | Vis-C<br>Def-B<br>TP<br>(N=3) | 250.6*<br>($p < 10^{-10}$) |
| Spatial Scale | Child | 550* | -0.1002* | Vis-C<br>Cont-A, Cont-B<br>Def-A, Def-B<br>(N=5) | Vis-P<br>Lim-A, Lim-B<br>Def-C<br>(N=4) | 300.6*<br>($p < 10^{-10}$) |
| | Adult | 701* | -0.1035* | Lim-B | Vis-C<br>SM-B | 61.4*<br>($p < 10^{-5}$) |
| | Old | 463* | -0.0449* | Lim-A, Limb-B<br>Cont-B<br>Def-C<br>(n=4) | Vis-C, Vis-P<br>SM-B<br>(N=3) | 255.7*<br>($p < 10^{-10}$) |
| Myelination | Child | 754* | 0.0097* | SM-A<br>DA-A, DA-B<br>Cont-C<br>(N=4) | SV-B<br>Lim-A, Lim-B<br>Def-A, Def-C<br>(N=5) | 456.9*<br>($p < 10^{-10}$) |
| | Adult | 905* | 0.0025* | (N=0) | (N=0) | 16.6<br>($p = 0.48$) |
| | Old | 566* | 0.0004* | SM-A<br>(N=1) | Lim-B<br>Cont-C<br>Def-C<br>(N=3) | 85.8*<br>($p < 10^{-10}$) |

To visualize the age-related changes in timescale within each age group, we fit the LOWESS curves to the timescales of individual participants for the entire cortex (Figure 3b) and also separately for different FC networks (Figure 3c). The average timescale for the entire brain as well as the timescales of individual networks tended to decrease with age. However, these plots also revealed non-monotonic age-related changes in the timescales, which was further tested using a regression model that included a quadratic age term. The average timescale for the entire brain showed a significant non-monotonic trend in the Child age group (p<0.005), reaching its peak at 11.1 years of age (Figure 3b). Significant non-monotonic age-related changes in timescales were also found for 13 of 17 FC networks, but only in the Child age group (Figure 3c). The peak age varied from 6 to 16 across these 13 networks, with the mean of 11.9 years of age (Figure 3d). This implies that the timescales of rs-fMRI signals still increase during early childhood, and do not reach their full maturation until adolescence. Therefore, while age has a global effect on timescales of rs-fMRI signals, these age-related changes vary across functional networks.

We found that throughout the lifespan, the timescales of the global signal were longer than the average timescales of rs-fMRI signals estimated with the global signal regression (paired t-test, t>80.0, p<10^−100^ for all age groups; Figure 3b and 4a). In addition, the timescale of the global signal significantly decreased with age in the Adult and Old age groups (p<0.01), although this change was significantly smaller than for the average timescale of FC networks estimated with the global signal regression (z-test, p<10^−6^). By contrast, for the Child age group, the timescale of the global signal showed a significant non-monotonicity, peaking at the age of 16.2. Thus, the timescales of the global signal and regional rs-fMRI signals showed qualitatively similar age-related changes. Nevertheless, we found that the timescales of rs-fMRI for individual FC networks were largely unaffected by the global signal regression. The timescales of rs-fMRI signals from different FC networks with and without the global signal regression were highly correlated in all age groups (r > 0.97, p < 0.001; Figure 4b).

**Figure 4.**
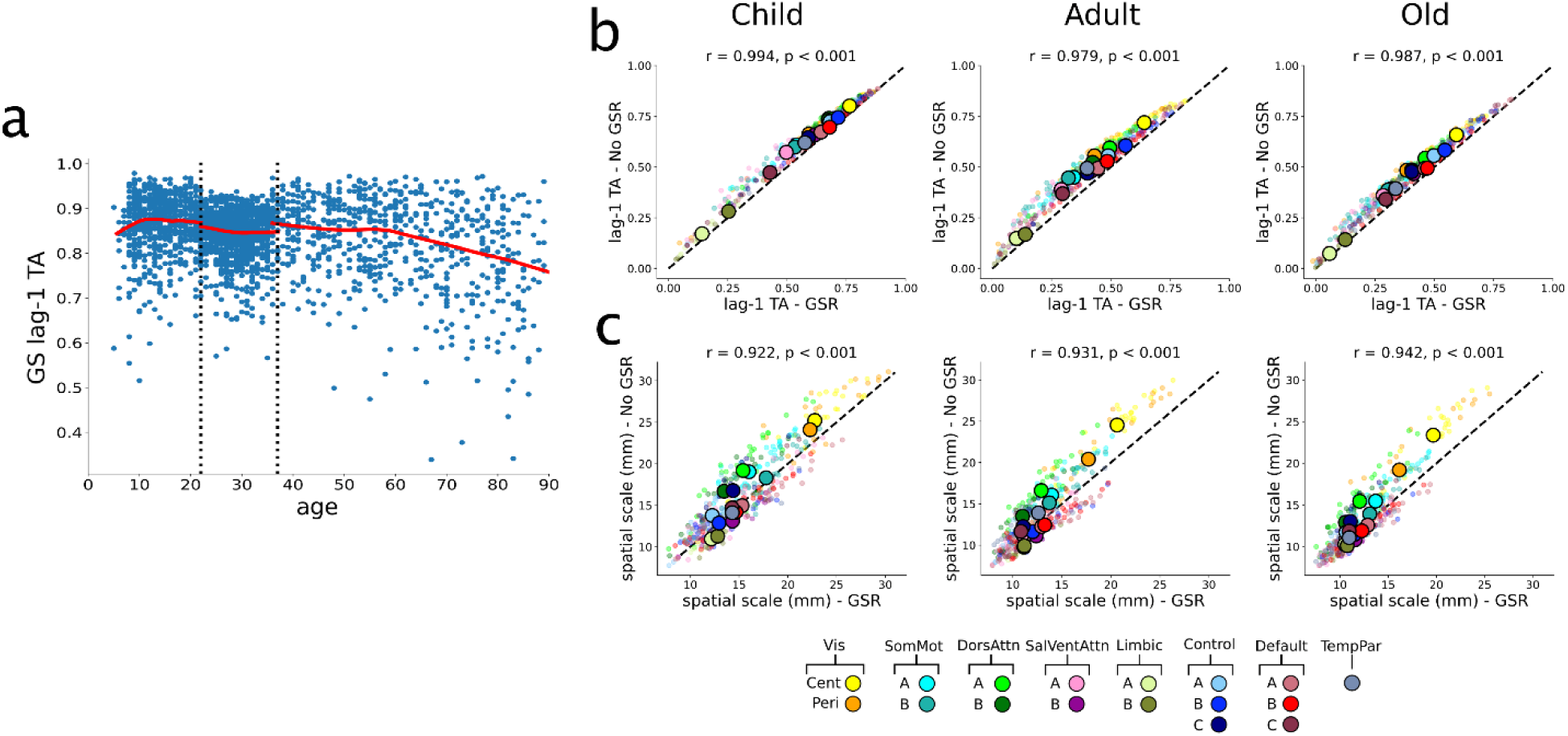
Age-related changes in the global signal and its effect on the temporal and spatial scales of rs-fMRI activity. (a) Timescale of the global signal across individual subjects. Red line indicates LOWESS fit. (b) and (c) Scatter plots for temporal (b) and spatial (c) scales estimated with (ordinate) and without (abscissa) the global signal regression.

### Age-related changes in spatial scales of fMRI signals

Next, we examined age-related changes in spatial scale of rs-fMRI signals. The brain map of the average spatial scale in each age group showed that the spatial scales tended to be larger in the somatosensory-motor, visual, and frontopolar regions (Figure 5a). These maps were robust, although the correlation coefficient between the randomly divided two halves was somewhat lower (r = 0.63, 0.60, and 0.58, for Child, Adult, and Old groups, respectively) than for the timescales. We also found strong and significant correlations between the maps of spatial scales across age groups (Corr > 0.95, p < 0.001; Figure 4b and 4c). Although the anatomical distribution of spatial scales remains similar across different age groups, the whole-brain average spatial scale decreased significantly and consistently across three different age groups (mean ± SD = 15.5 ± 6.0, 13.1 ± 5.1, and 12.5 ± 5.0 mm for Child, Adult, and Old groups, respectively; 1-way ANOVA, p<0.05).

**Figure 5.**
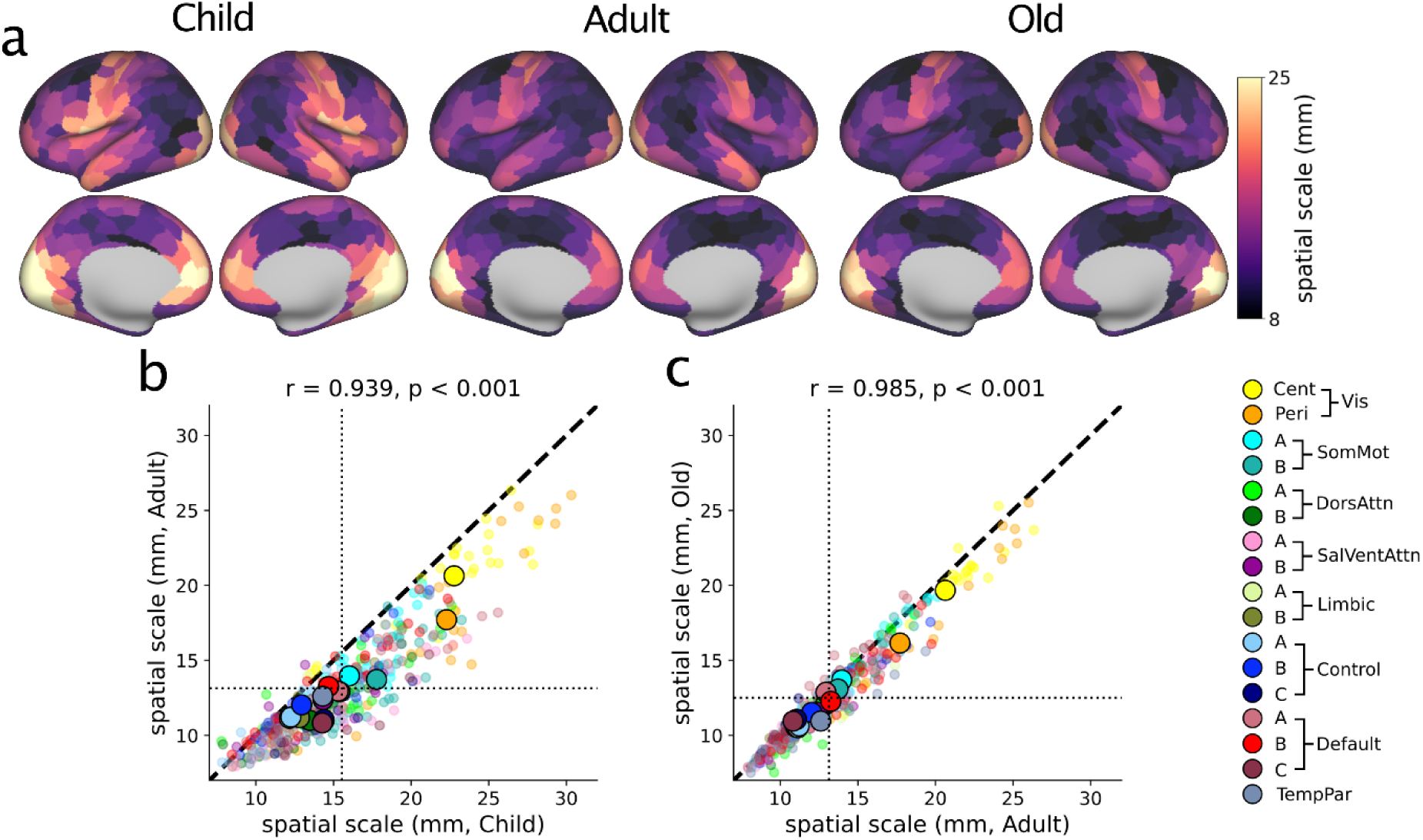
Spatial scales of rs-fMRI activity. (a) Brain maps of mean spatial scale of rs-fMRI signals for Child (left), Adult (center), and Old (right) age groups. (b) and (c) Scatter plots showing spatial scales for Child vs Adult (b) and Adult vs Old (c). Dotted lines show group means.

Using linear mixed effects models, we also found that spatial scales decreased with age in all age groups (Table 2; Figure 5a). This analysis also revealed significant age × network interactions for some networks in each age group (n=9, 3, and 7 for Child, Adult, and Old groups, respectively; Table 2). Similar to the pattern with the timescale, some FC networks show significant age × network interactions in both Adult and Old age groups. For example, Limbic-B network showed significantly less negative slopes than the average in both age groups, whereas Visual-Cent and Somatosensory-Motor-B networks showed significantly more negative slopes (Table 2).

The age coefficients for spatial scales in individual ROI showed that spatial scales tended to decrease with age for most brain areas throughout the lifespan. However, similar to the age coefficients for timescales, they were significantly positive for some areas of the prefrontal cortex in the Child age group (Figure 6a). Furthermore, although LOWESS fits to the whole brain and network averages for subjects in each group similarly showed overall decrease in spatial scales with age (Figure 6b), some networks displayed non-monotonic trajectories in the Child age group (Figure 6c). Although the number of FC networks for which the quadratic regression model fit age-related changes in spatial scale better was smaller (n=5) than that for the timescales (n=13), the average peak ages were similar (mean=11.9 and 11.3 for temporal and spatial scales, respectively; Figures 3d and 6d). Similar to the timescales of rs-fMRI signals, we also found that the spatial scales estimated with and without the global signal regression were highly correlated (r>0.9, p<0.001; Figure 4c).

**Figure 6.**
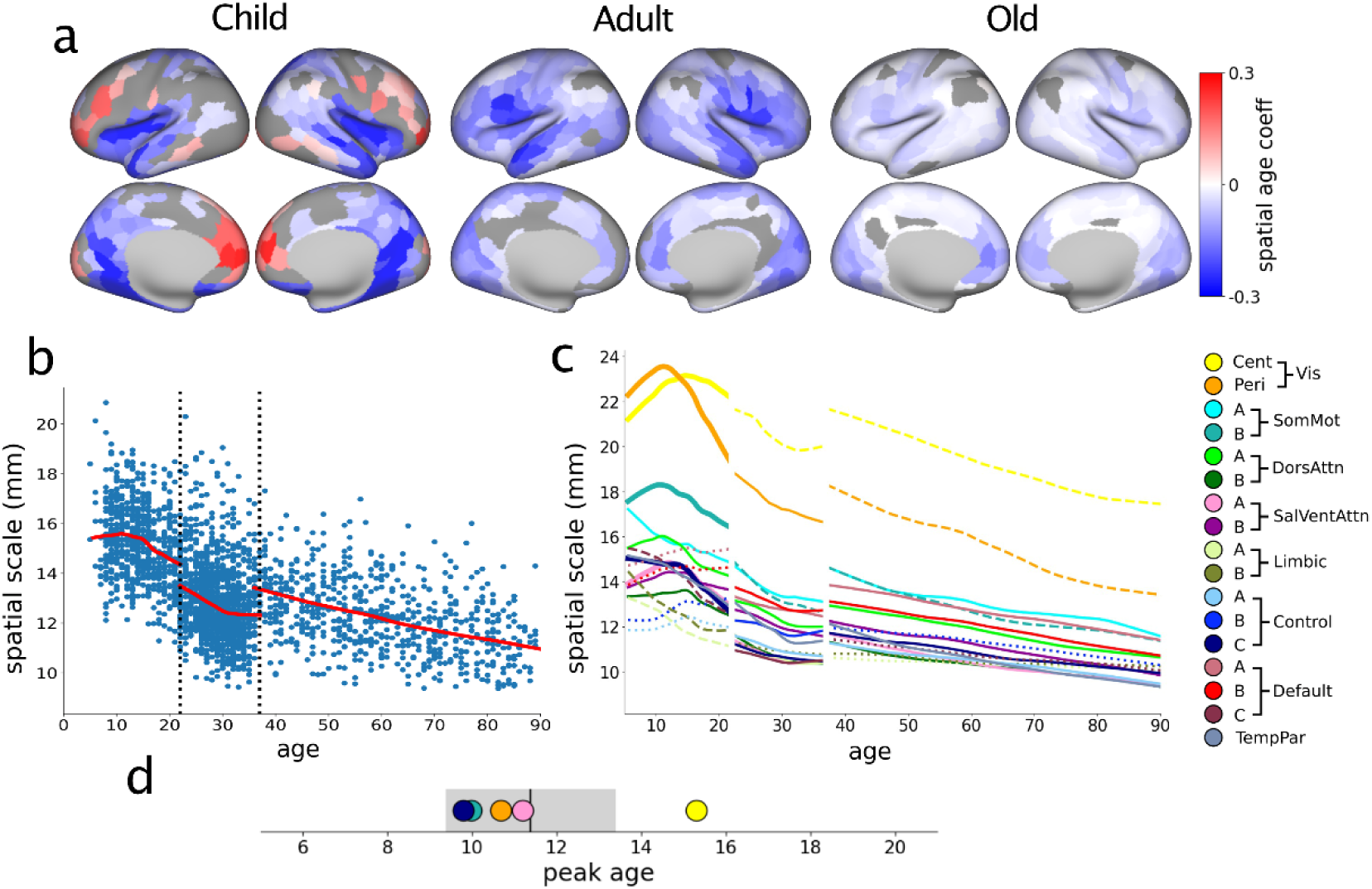
Age-related changes in the spatial scale of rs-fMRI signals. (a) Brain maps of the age coefficients for the spatial scale (SA-λ) in Child (left), Adult (center), and Old (right) age groups. Gray regions indicate the ROI in which the age effect was not statistically significant (p>0.05). (b) Scatter plots of the average spatial scale for the entire cortex across individual subjects. (c) LOWESS fit for the spatial scale in each FC network. Dotted/dashed lines show networks that are significantly more positive/negative than the average. (d) Peak ages for spatial scale in networks with significant non-monotonic timecourse in the Child age group. Same format as in Figure 3.

### Comparison of temporal and spatial scales

The above results suggest that age-related changes in temporal and spatial scales of fMRI signals might show similar patterns. In particular, the age-related changes in temporal and spatial scales were non-monotonic for some prefrontal regions in the Child age group, which would not be fully captured by the age coefficient in a linear regression model. Therefore, for each network and age group, we tested whether these two scales were correlated across different subjects. We found that these two scales were strongly and significantly correlated for all networks in each age group (mean correlation coefficient ± SD = 0.64 ± 0.09, 0.74 ± 0.10, 0.74 ± 0.09, for Child, Adult, and Old age groups, respectively). The correlations between the two scales for the networks with non-monotonic trajectories in the Child age group ranged from 0.45 to 0.78, and they were not significantly different from the monotonic network correlations (t-test, p>0.05). We found that the correlation between the temporal and spatial scales at the network level was not statistically significant (Figure 7, top). On the other hand, we found a significant correlation between the age coefficients for temporal and spatial scales in each age group (Figure 7, bottom), suggesting that the age-related changes in temporal and spatial scales of rs-fMRI signals might be coordinated throughout the lifespan.

**Figure 7.**
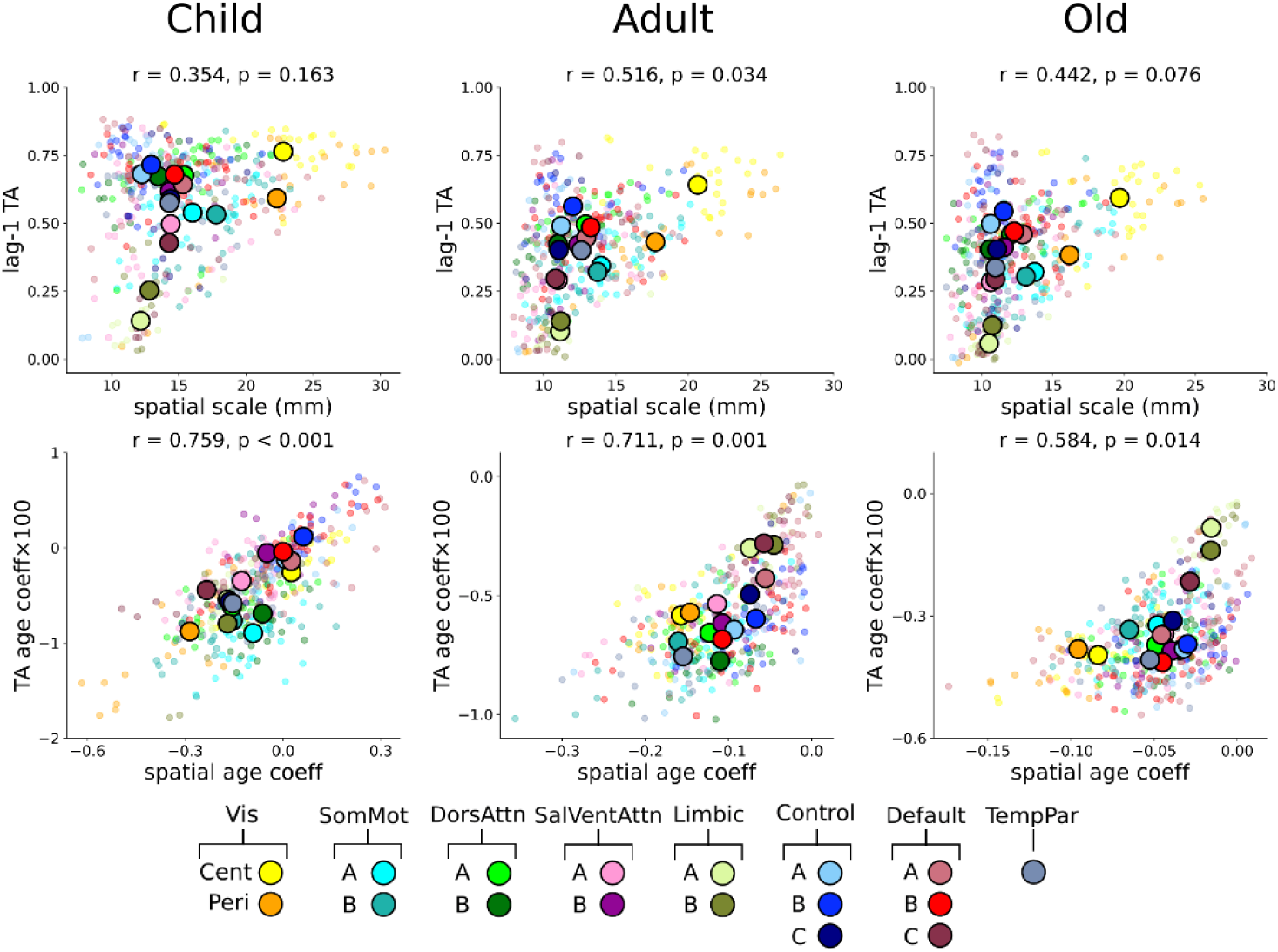
Relationship between the temporal and spatial scales of rs-fMRI activity. Scatter plots comparing the temporal and spatial scales (top) and the corresponding age coefficients (bottom). Small transparent dots indicate the values for individual ROI, while the large opaque circles indicate network averages.

Next, we examined further whether some FC networks tend to show non-monotonic age-related changes consistently for temporal and spatial scales. Given that the results in the present study were based on 3 separate datasets, non-monotonic patterns manifest in two different ways. First, the results from quadratic regression models have shown that for some FC networks in the Child age group, the temporal and spatial scales tend to peak approximately at the age of 12 (Figure 8 left, green). Second, we also found that for some prefrontal areas in the Child age group, the age coefficients for both scales were significantly positive (Figure 8 left, red). Given that the age coefficients for both scales were significantly negative for all areas in the Adult age group (22 years and older), this suggests that the temporal or spatial scale might reach their peak values around the age of 22 for areas with positive age coefficients in the Child age group. Therefore, we looked for the ROI that showed either a significant non-monotonic trend or a significant positive age coefficient in both temporal and spatial scales. For example, some areas showed positive age coefficient for the spatial scale but a quadratic change for timescale in the Child age group (Figure 8 right, yellow). The results from this analysis revealed that for some regions in the prefrontal and parietal cortical areas as well as the insular cortex, both temporal and spatial scales might increase and reach their peak values before the age of 22 (Figure 8 right).

**Figure 8.**
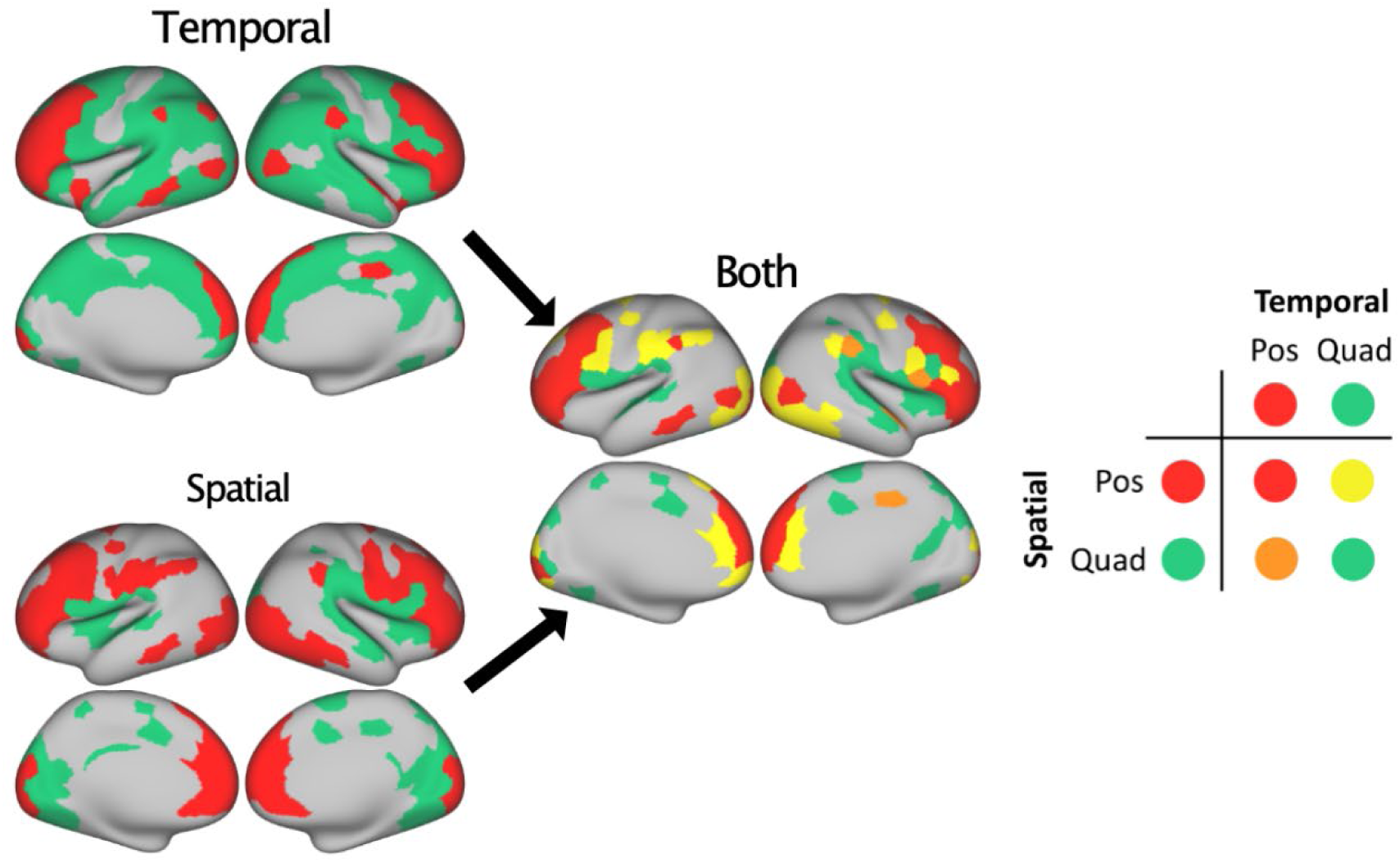
Brain maps showing non-monotonic age-related changes in temporal and spatial scales. The ROI with significant positive age coefficient in the Child age group are shown in red, while the regions with a significant quadratic trend in the same group are shown in green for temporal (top left) and spatial (bottom left) scales. The ROI showing either significant quadratic trend or significantly positive age coefficient for both scales are also shown (right) with different colors indicating different combinations of these two patterns.

### Age-related changes in cortical myelination

Next, we examined the regional variability in cortical myelination and its age-related changes. Consistent with the results in previous studies (Grydeland et al., 2019), brain maps of average myelin show an overall increase from childhood to adulthood and became stable in the Old age group, with the highest values seen around the central sulcus and posterior medial regions near the calcarine fissure (Figure 9a). Similar to the brain maps for temporal and spatial scales, these myelin maps were highly correlated across different age groups at the network level (r > 0.85, p < 0.001; Figure 9b and 9c).

**Figure 9.**
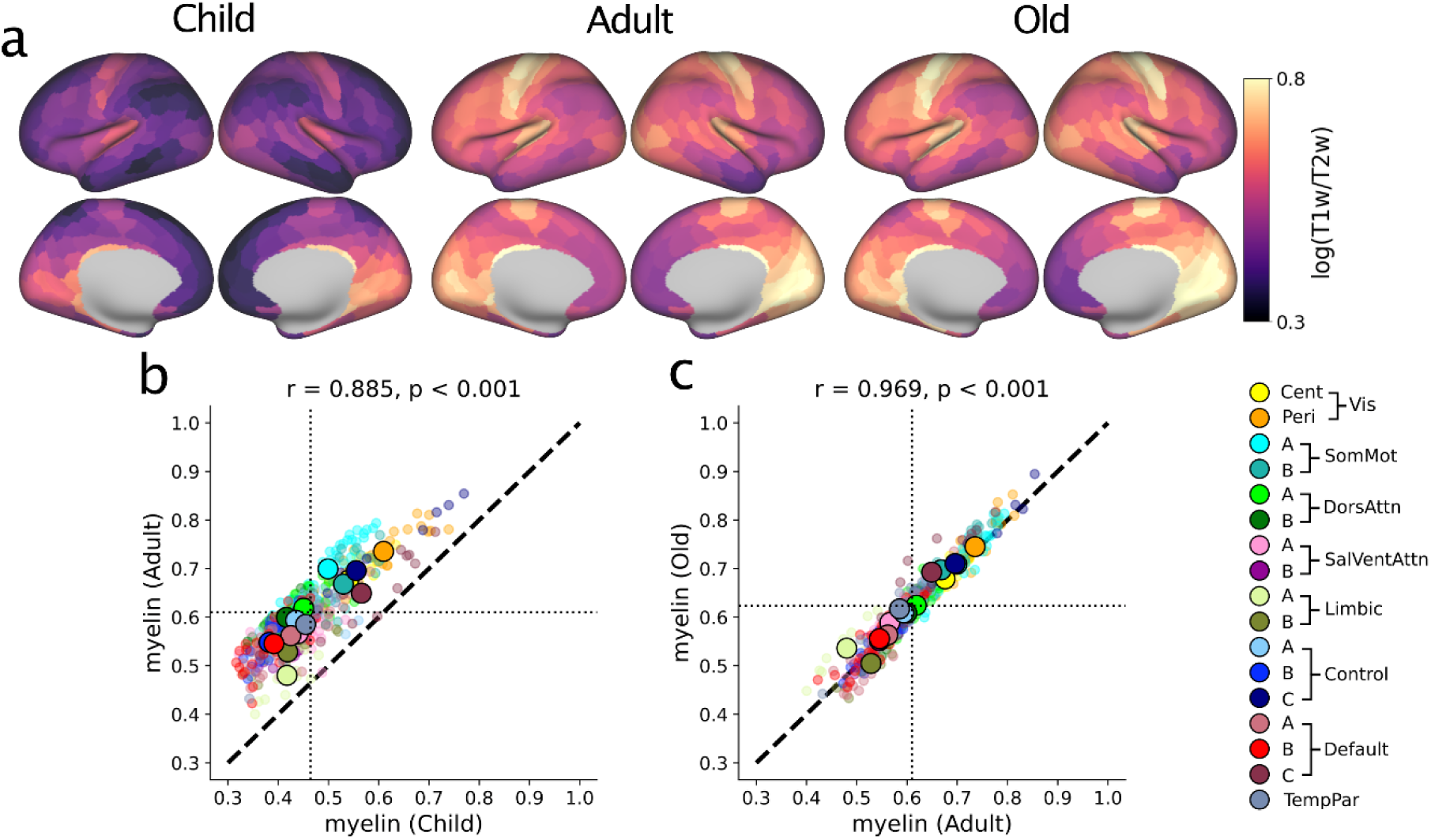
Cortical myelination across the lifespan. (a) Brain maps of the mean myelination maps for Child (left), Adult (center), and Old (right) age groups. (b) and (c) Scatter plots for comparing the mean myelinations for Child vs Adult (b) and Adult vs Old (c) age groups. Dotted lines show group means.

Cortical myelination increased steeply throughout the brain in the Child age group and more slowly in the Adult and Old age groups (Figure 10a). In particular, the age coefficient for cortical myelination was positive in all FC networks for the Child age group, and a significant non-monotonic trend was not observed either for the brain average (Figure 10b) or for any individual networks (Figure 10c). In addition, the full linear mixed effects model did not fit the age-related changes in cortical myelination better than the model without the interaction terms in the Adult age group (Table 2), indicating that cortical myelination changes uniformly and consistently throughout the brain during adulthood. By contrast, the full model fit the results better in the Child and Old age group, and the rate of age-related changes for cortical myelination was significantly different from the mean for some FC networks. For example, Somatosensory-Motor-B network showed a significantly more positive slope than the average in both Child and Old age groups, whereas Limbic-B and Default-C networks showed significantly more negative slopes in both age groups (Table 2).

**Figure 10.**
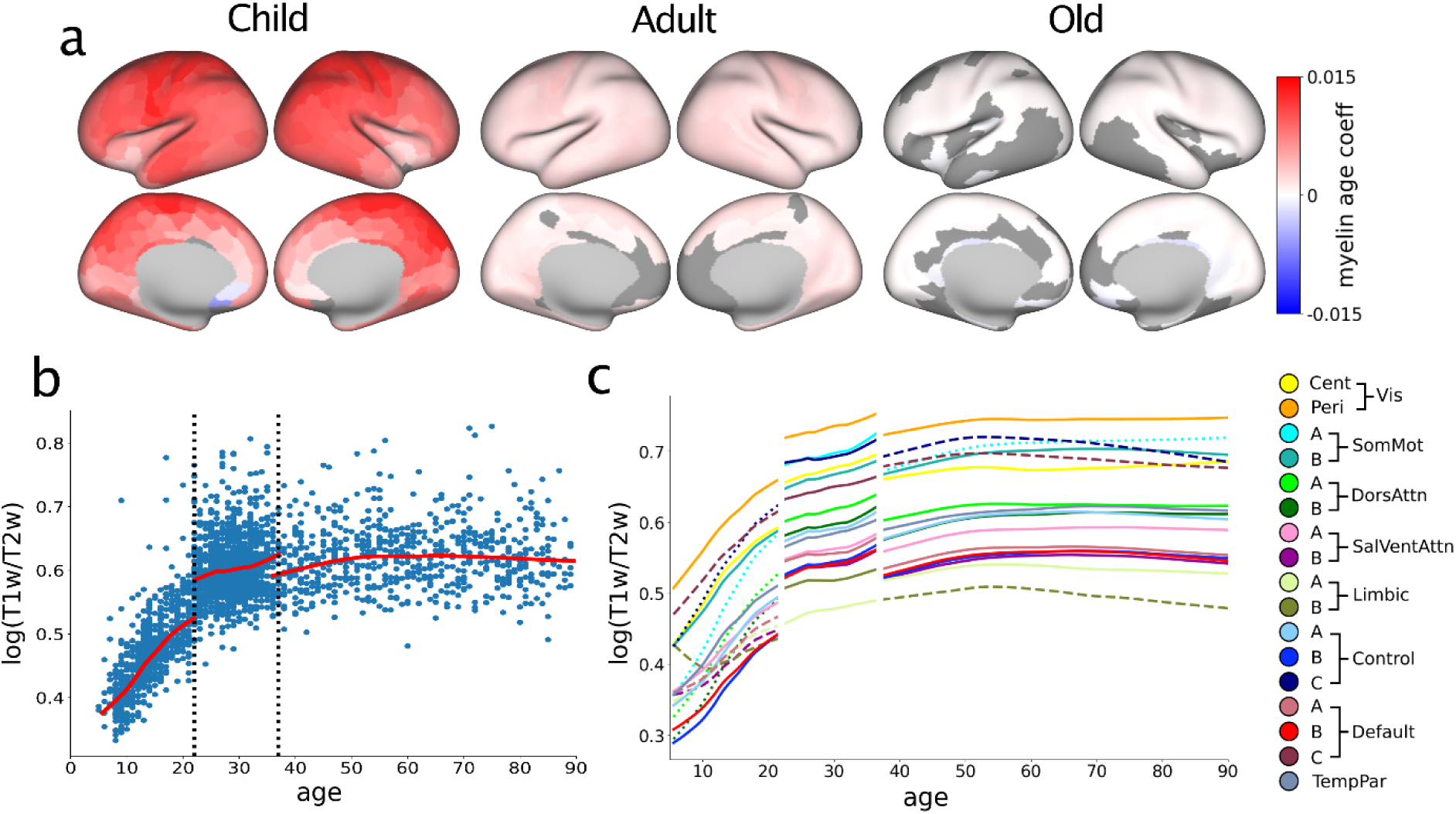
Age-related changes in cortical myelination. (a) Brain maps of the age coefficient for myelination in Child (left), Adult (center), and Old (right) age groups. Gray regions indicate the ROI in which the age effect was not statistically significant (p>0.05). (b) Scatter plots of the average myelination for the entire cortex across individual subjects. (c) LOWESS fit for myelination in each FC network. Dotted/dashed lines show networks that are significantly more positive/negative than the average.

We then examined whether and how cortical myelination and its age-related changes might be related to temporal and spatial scales of rs-fMRI activity across different age groups. First, we found that cortical myelination was not significantly correlated with the timescale of rs-fMRI signals across FC networks in any age group (r = 0.02, 0.32, 0.21 for Child, Adult, and Old age groups, respectively, p>0.2; Figure 11 top). Second, the age coefficients for timescale and myelination were significantly correlated for the Adult (r = −0.51, p<0.05) and Old age groups (r = −0.66, p<0.005), but not for the Child age group (r = −0.182, p>0.4; Figure 11 bottom). This suggests that the age-related changes in temporal scale and myelination are more closely coordinated during adulthood than during childhood. Third, we found that spatial scale of rs-fMRI activity was significantly correlated with cortical myelination across FC networks in all age groups (r = 0.70, 0.53, 0.50 for Child, Adult, and Old age groups, respectively, p<0.05; Figure 12 top). Finally, the age coefficients for spatial scale and cortical myelination were significantly correlated only in the Adult age group (r = 0.15, −0.64, and −0.41 for the Child, Adult, and Old age groups, respectively; Figure 12 bottom).

**Figure 11.**
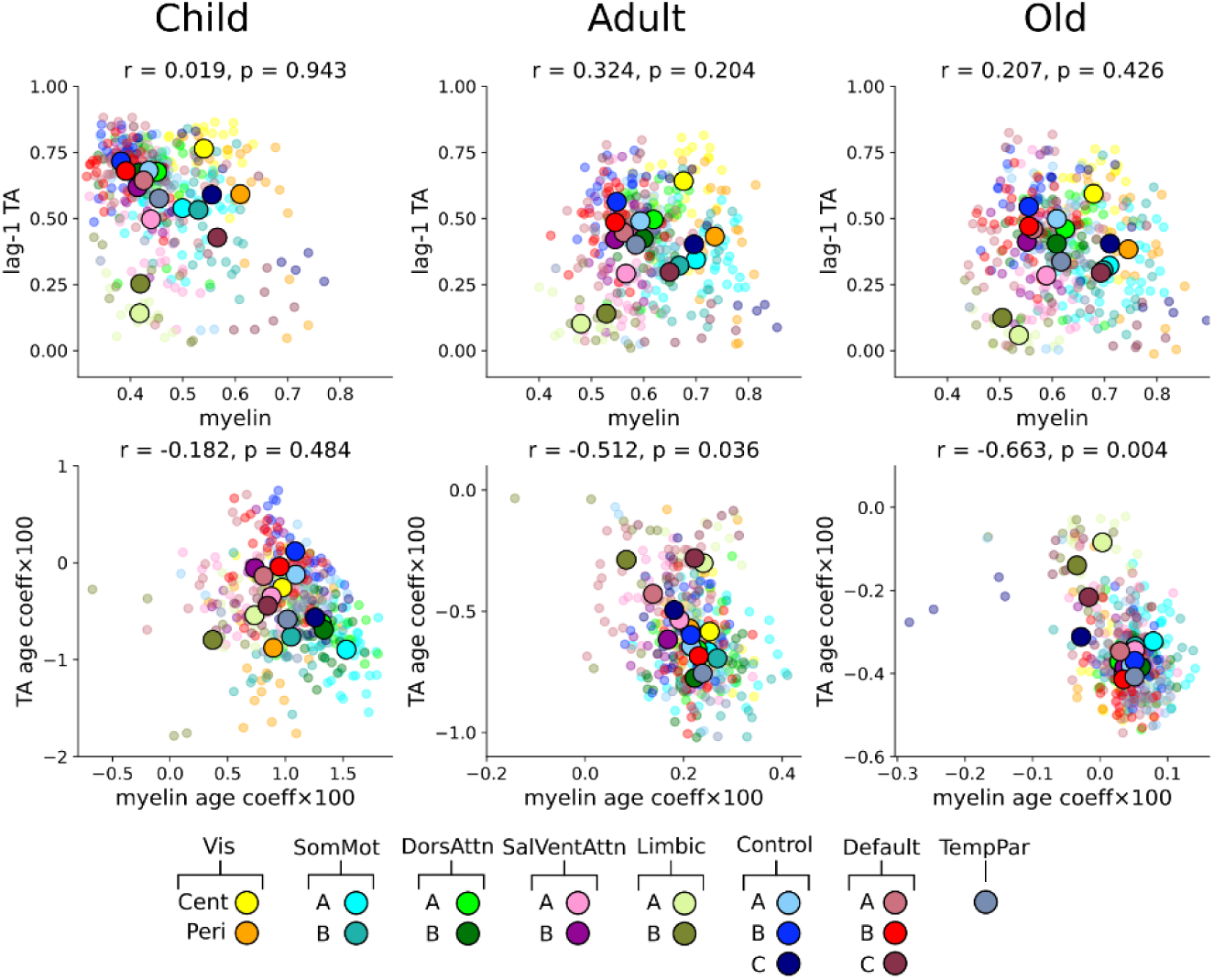
Relationship between timescale of rs-fMRI activity and myelination. Scatter plots for cortical timescale of rs-fMRI signals and cortical myelination (top) and their corresponding age coefficients (bottom). Small transparent dots indicate the values for individual ROI, while the large opaque circles indicate network averages.

**Figure 12.**
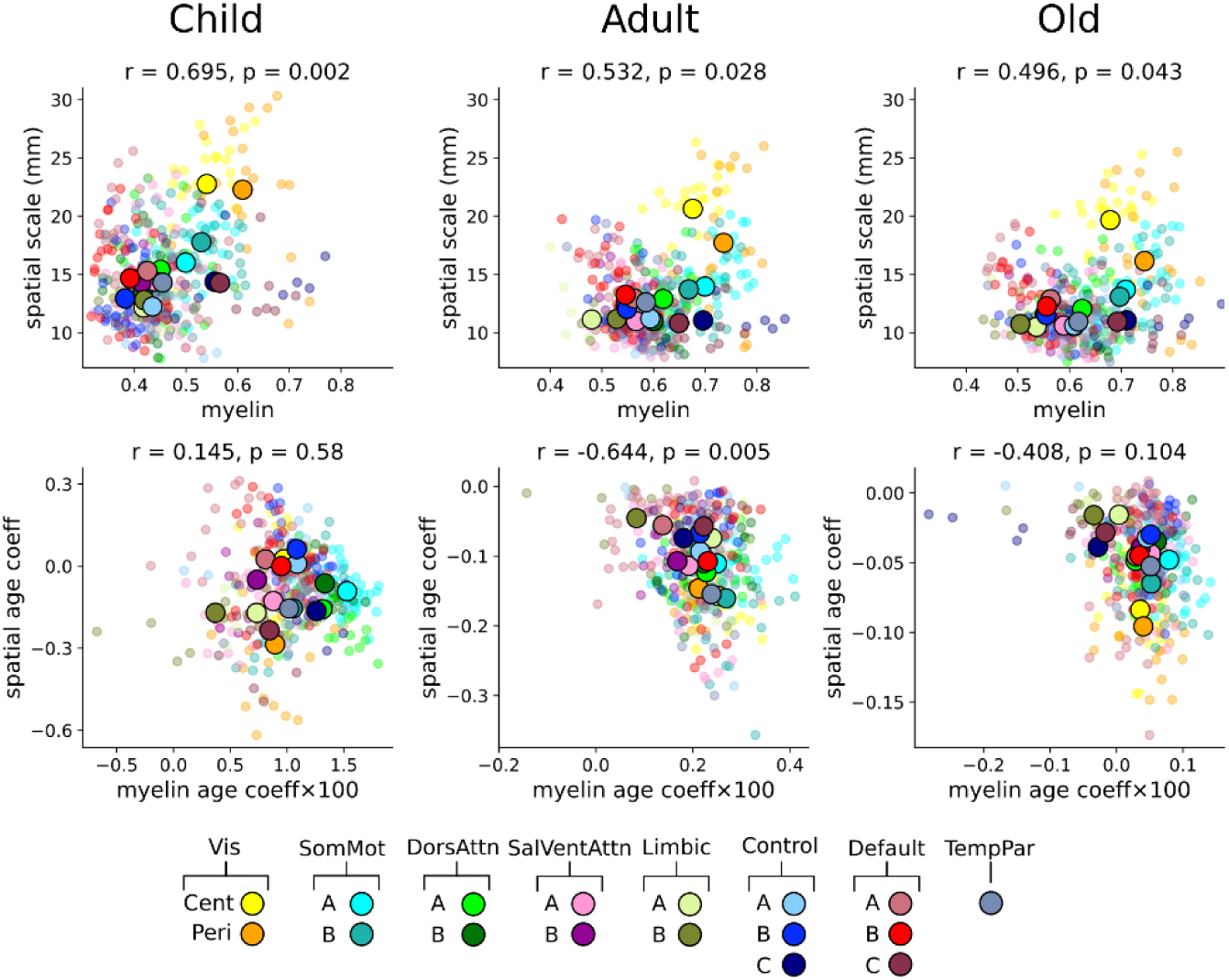
Relationship between spatial scale of rs-fMRI activity and myelination. Scatter plots for spatial scale of rs-fMRI signals and cortical myelination (top) and their corresponding age coefficients (bottom). Same format as in Figure 11.

### Relationship among multiple cortical gradients

We compared the brain maps for temporal and spatial scales of rs-fMRI activity and cortical myelination together with their corresponding age coefficients to the ranking along the sensorimotor-association (S-A) axis (Figure 1b; Figure 13). Consistent with the findings from previous studies (Sydnor et al., 2023), we found that the cortical myelination had a strong and significant negative correlation with the ranking along the S-A axis (Figure 13, right). We also found a significant negative correlation between the spatial scale and the rank along the S-A axis at both ROI and network levels in each age group, indicating that sensory cortical areas tended to have longer spatial scales (Figure 13, center). We did not find any significant correlation between the timescale of resting-state activity and the rank along the sensorimotor-association axis in any age group (Figure 13, left). On the other hand, we found that the age coefficients for the timescale were significantly correlated only for the Child age group at both ROI and network levels.

**Figure 13.**
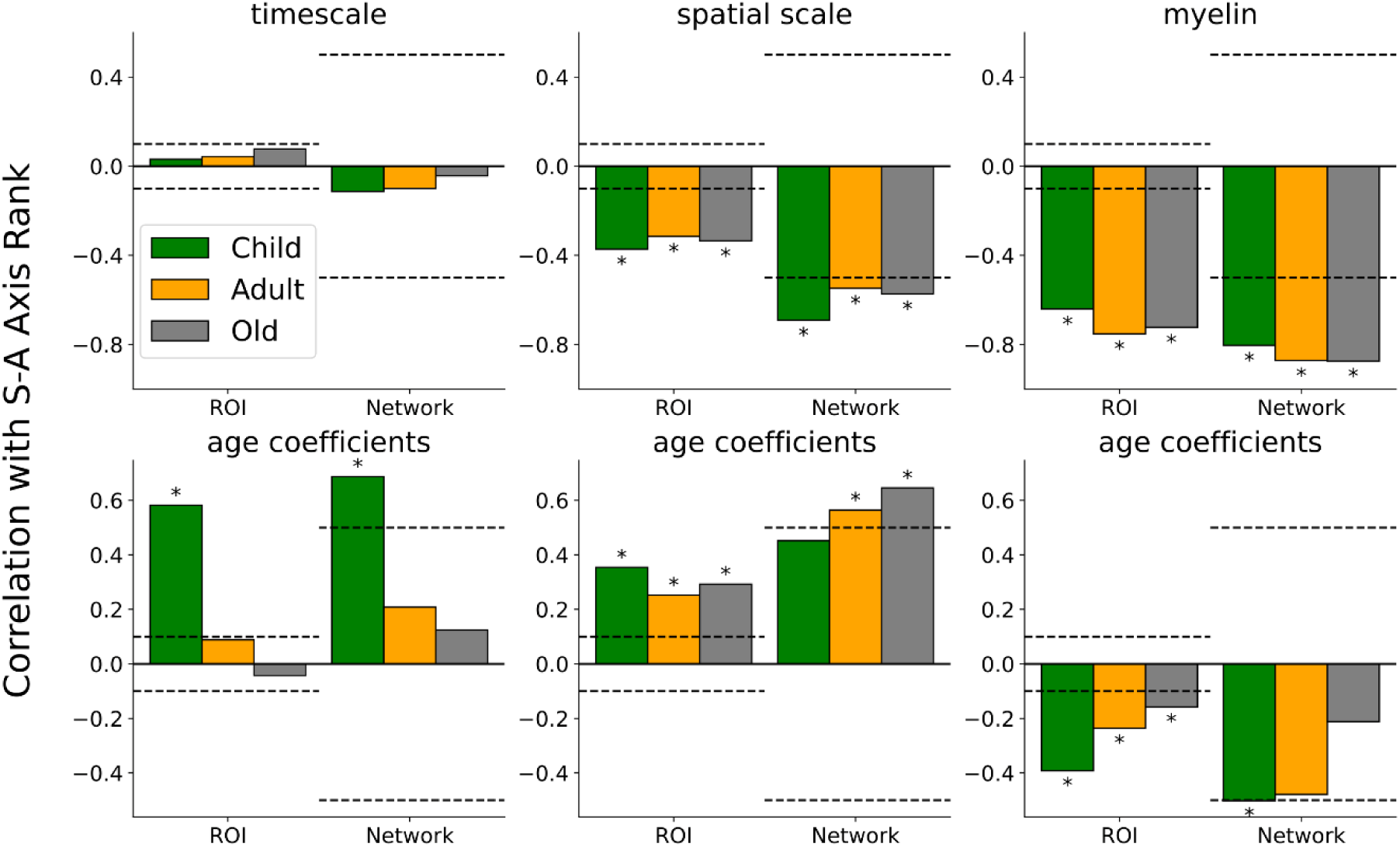
Relationship between the ranking along sensorimotor-association (S-A) axis and three measures examined in the present study (timescale, spatial scale, and cortical myelination; top) and their corresponding age coefficients (bottom) at ROI and network levels. Significance thresholds are indicated by dashed lines, with an asterisk indicating significant correlations for a given age group (p<0.05).

To examine more systematically how the similarities among these different maps change across different age groups, we visualized the correlations among various brain maps in each age group as a graph in which an edge or link between any two nodes represents the correlation between the two corresponding brain maps (Figure 14). These graphs show that some brain maps consistently show significant correlations across all age groups. For example, spatial scale and myelination were positively correlated, whereas both spatial scale and cortical myelination were negatively correlated with the S-A axis ranking, in all age groups (Figure 14, light colors). In addition, correlations between some of these measurements were significant only in some age groups (Figure 14, dark colors). For example, the negative correlation between the age coefficient for the spatial scale and the cortical myelination was present only in the Child and Old age groups, suggesting that they might be coordinated only during development and aging. On the other hand, correlations between some brain maps were significantly only in the Child age group, suggesting that their age-related changes might be co-regulated during development. For example, the age coefficient for the timescale was positively and negatively correlated with S-A rank and cortical myelination only for the Child age group. Finally, coupling between some measures appeared only in the Adult and Old age groups, such as the negative correlation between the age coefficients for timescale and cortical myelination, and the positive correlation between the age coefficient for spatial scale and S-A rank.

**Figure 14.**
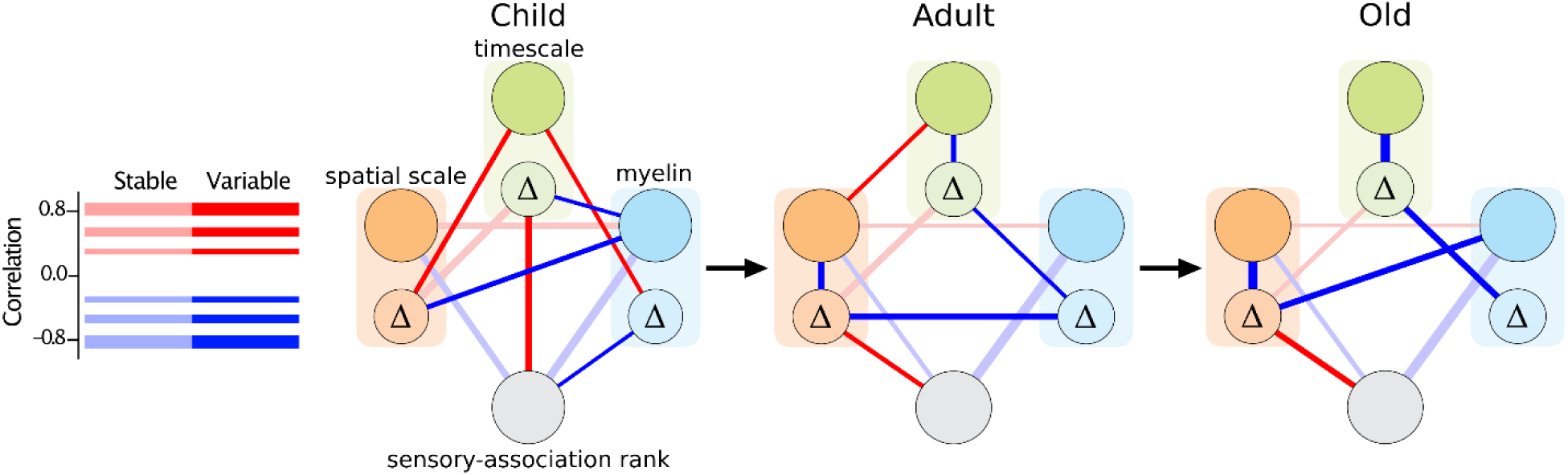
Relationship between multiple cortical maps for different age groups. Correlations among different brain maps for temporal and spatial scales of rs-fMRI activity, cortical myelination, and the ranking along the S-A axis are visualized as an edge between the two nodes corresponding to different brain maps. The brain maps for the age coefficients for timescale, spatial scale, and myelination are indicated by the circles with symbol Δ. Only connections between the maps with significant correlations are shown, with the color and thickness of the line indicating the sign and magnitude of correlation coefficient. Significant correlations consistent across all age groups are shown in light colors.

## Discussion

In the present study, we found that temporal and spatial scales of resting-state fMRI signals in the human cortex tended to decrease with age, while they increased in some prefrontal regions during childhood. Although the regional variations in these two scales were not correlated consistently across different age groups, the rates of their age-related changes were correlated across cortical networks in all age groups. We also found that the age-related changes in spatial and temporal scales were closely coordinated with the level of cortical myelination or the rate of its age-related changes across the lifespan. Although we did not find any correlation between the timescale and the ranking along the sensorimotor-association (S-A) axis, the rate of age-related changes in the timescale increased along the S-A axis in the Child age group. Finally, both the spatial scale and myelination decreased along the S-A axis consistently in all age groups, whereas the rate of age-related changes in the spatial scale increased along the S-A axis in the Adult and Old age groups.

### Temporal and spatial scales of resting-state activity

In previous studies, the so-called intrinsic timescale of cortical activity was often measured by the decay rate of the temporal autocorrelation in neural activity (Murray et al., 2014; Ito et al., 2020; Raut et al., 2020). Here, we used the lag-1 autocorrelation to measure the timescale in rs-fMRI activity for several reasons. First, the lag-1 autocorrelation is strongly correlated with the first principal component extracted from diverse temporal features of the resting-state activity (Shafiei et al., 2020). Second, the regional variation in the lag-1 autocorrelation reliably captures the individual variability in the resting-state activity and its graphic topology (Shinn et al., 2023). Previous studies (Demirtaş et al., 2019, Yeo et al., 2011), especially those based on electrophysiological recordings (Murray et al., 2014; Gao et al., 2020; Wolff et al., 2022; Song et al., 2024; Siegle et al., 2021; Manea et al., 2022; Manea et al., 2024), have found a hierarchical ordering of timescales across the cortex, with unimodal and transmodal areas exhibiting shorter and longer timescales, respectively (Murray et al., 2014; Ito et al., 2020; Raut et al., 2020). By contrast, we found that the timescale estimate based on the lag-1 autocorrelation was not significantly related to the hierarchical gradient along the sensorimotor-association axis in any of the age groups (see also Shinn et al., 2023). In fact, a recent study also found that the first principal component estimated from various temporal features of rs-fMRI activity does not follow such a gradient (Shafiei et al., 2020). These results suggest that the resting-state BOLD activity might be characterized by multiple timescales (Spitmann et al., 2020; Song et al., 2024). By contrast, using a novel method to estimate the anatomical gradient of the spatial scale in the rs-fMRI activity, we found that spatial scale was significantly related to the ranking along the S-A axis.

We also found that temporal and spatial scales of rs-fMRI activity undergo systematic changes in coordination with cortical myelination. Sensory and motor cortical areas are more myelinated than association cortical areas (Flechsig, 1901; Glasser and Van Essen, 2011; Huntenburg, 2018). This might reflect different levels of plasticity across cortical areas, since myelination inhibits neural plasticity (Takesian and Hensch, 2013). Previous studies showed that the time course of myelination as well as the level of myelination differs along the sensory-association cortical axis, proceeding earlier in the unimodal sensory and motor areas than the transmodal association cortical areas (Grydeland et al., 2019; Baum et al., 2022). Consistent with these previous findings, we found that the rate of myelination in the Child age group was negatively associated with the ranking along the S-A axis. In addition, the rate of age-related changes in the timescale during childhood was positively correlated with the ranking along the S-A axis during childhood and negatively with the changes in cortical myelination during adulthood and aging. This suggests that the factors regulating the timescale of rs-fMRI activity might differ for development and aging. By contrast, we found that the spatial scale was related to myelination and S-A axis consistently throughout the lifespan, although it decreased faster in sensory and motor cortical areas than in association cortical areas during adulthood and aging. In addition, during adulthood, age-related changes in spatial scale and cortical myelination were correlated. These results suggest that throughout the lifespan age-related changes in temporal and spatial scales might be influenced differently by various anatomical and functional properties of cortical networks. Timescales of resting-state fMRI may be altered in some clinical conditions (Zhang et al., 2024; Uscătescu et al., 2023) and related to memory performance (Wu and Gollo, 2025). However, more studies are needed to understand the specific aspects of cognitive functions that can be predicted by the temporal and spatial scales of resting-state activity.

### Implications for brain development and aging

Several studies have found an age-related decrease in intrinsic timescales of BOLD (Geerligs et al., 2017; Truzzi and Cusack, 2023; Wu and Gollo, 2025) and EEG (Gao et al., 2020) activity in the human cortex, although they did not cover the entire lifespan. Consistent with these previous findings, we found that the timescale of rs-fMRI activity mostly decrease throughout the lifespan. We also found that the spatial scale of rs-fMRI activity decreased with age. Given that the resting-state functional connectivity is dominated by the relatively low-frequency power (Cordes et al., 2001), an age-dependent decrease in the spatial scale of BOLD activity is consistent with the previous findings that the strength of functional connectivity tends to decrease with age (Ferreira and Busatto, 2013). Age-related changes in the temporal and spatial scales of rs-fMRI activity also suggest that functions of spontaneous activity related to plasticity might change throughout the lifespan. Longer spatial and temporal scales in spontaneous activity early during development might allow developing brains to integrate incoming information from their surroundings over longer spatial and temporal windows, helping them to learn more general features (Kiebel et al., 2008; Vogelstang et al., 2018). Subsequent decorrelation and reduced synchrony in neural activity during later development might contribute to finer encoding of temporal and spatial information (Golshani et al., 2009; Gribizis et al., 2019; Martini et al., 2021).

We found that age-related changes in temporal and spatial scales were positively correlated in every age group, indicating that changes in both scales might be coordinated throughout the lifespan. On the other hand, a recent study showed the spatial scale estimated globally for the entire brain increased with age (Shinn et al., 2023). This discrepancy is likely due to the differential effects of age on local vs. global functional connectivity, which has been attributed to delayed maturation of higher-order cognitive systems in the brain (Betzel et al., 2014; Deery et al., 2023; Supekar et al., 2010). Consistent with an age-related decline in the low-frequency fluctuation in the global signal reported previously (Ao et al., 2022), we found that the timescale of the global signal tended to decrease with age. Age-related changes in both temporal and spatial scales of rs-fMRI activity were, however, unchanged by whether the global signal was factored out or not, suggesting that they might arise from different underlying mechanisms.

For some regions in the prefrontal cortex, we found that both temporal and spatial scales increased with age during childhood and peaked between the ages of 11 and 12. Previous studies showed that the age-related changes in the average power of low-frequency (<0.1 Hz) fluctuation in the resting state BOLD signals decreased monotonically in sensory and motor areas but increased in the prefrontal cortex during childhood and started to decrease late in adolescence (Sydnor et al., 2023). The finding that functional connectivity in the default mode network changed non-monotonically during childhood (Faghiri et al., 2019) is also consistent with the changes in the timescale observed in the present study. These results provide additional support to the hypothesis that maturation of the prefrontal cortex might be more delayed compared to the rest of the brain (Fransson et al., 2011; Dumontheil et al., 2008; Sydnor et al., 2021).

Some of the age-related changes in temporal and spatial scales reported in this study might be due to changes in hemodynamic response. In fact, previous studies have found age-related changes in hemodynamic response functions during development (Arichi et al., 2012; Schmithorst et al., 2015) and aging (Abdelkarim et al., 2019; West et al., 2019; Tsvetanov et al., 2021; Turner et al., 2023). Nevertheless, the exact contribution of age-related changes in neurovascular coupling to timescales of resting-state activity is still not fully understood. This is difficult to study because both spontaneous activity (Khazipov and Luhmann, 2006) and neural response to sensory stimuli change during development and aging (Lippé et al., 2007; Fabiani et al., 2014; Gao et al., 2020; Chettouf et al., 2022; Springer et al., 2023). More research is therefore needed to elucidate the precise manner in which the temporal and spatial changes in BOLD activity reflect the neural and neurovascular factors.

### Software Accessibility

The final numerical values used to make all figures in the current study, along with the code to generate them, are available from the authors upon request.

## Acknowledgments

Data used in the preparation of this manuscript were obtained from the National Institute of Mental Health (NIMH) Data Archive (NDA). NDA is a collaborative informatics system created by the National Institutes of Health to provide a national resource to support and accelerate research in mental health. Dataset identifier: 10.15154/5f0d-z111. This manuscript reflects the views of the authors and may not reflect the opinions or views of the NIH or of the Submitters submitting original data to NDA.

## References

Abdelkarim D, Zhao Y, Turner MP, Sivakolundu DK, Lu H, Rypma B (2019) A neural-vascular complex of age-related changes in the human brain: Anatomy, physiology, and implications for neurocognitive aging. Neurosci Biobehav Rev 107:927–944.

Achard S, Bassett DS, Meyer-Lindenberg A. Bullmore E (2008) Fractal connectivity of long-memory networks. Phys Rev E 77:036104.

Achard S and Gannaz I (2015) Multivariate wavelet Whittle estimation in long-range dependence. J Time Ser Anal 37:476–512.

Ao Y, Kou J, Yang C, Wang Y, Huang L, Jing X, Cui Q, Cai X, Chen J (2022) The temporal dedifferentiation of global brain signal fluctuations during human brain ageing. Sci Rep 12:3616.

Arbabshirani MR, Preda A, Vaidya JG, Potkin SG, Pearlson G, Voyvodic J, Mathalon D, Van Erp T, Michael A, Kiehl KA, Turner JA, Calhoun VD (2019) Autoconnectivity: A new perspective on human brain function. J Neurosci Methods 323:68–76.

Arichi T, Fagiolo G, Varela M, Melendez-Calderon A, Allievi A, Merchant N, Tusor N, Counsell SJ, Burdet E, Beckmann CF, Edwards AD (2012) Development of BOLD signal hemodynamic responses in the human brain. NeuroImage 63:663–673.

Baum GL, Flournoy JC, Glasser MF, Harms MP, Mair P, Sanders AFP, Barch DM, Buckner RL, Bookheimer S, Dapretto M, Smith S, Thomas KM, Yacoub E, Van Essen DC, Somerville LH (2022) Graded Variation in T1w/T2w Ratio during Adolescence: Measurement, Caveats, and Implications for Development of Cortical Myelin. J Neurosci 42:5681–5694.

Bero J, Li Y, Kumar A, Humphries C, Nag S, Lee H, Ahn WY, Hahn S, Constable RT, Kim H, Lee D (2023) Coordinated anatomical and functional variability in the human brain during adolescence. Hum Brain Mapp 44:1767–1778.

Bethlehem RAI et al. (2022) Brain charts for the human lifespan. Nature 604:525–533.

Betzel RF, Byrge L, He Y, Goñi J, Zuo X-N, Sporns O (2014) Changes in structural and functional connectivity among resting-state networks across the human lifespan. NeuroImage 102:345–357.

Blinkouskaya Y, Caçoilo A, Gollamudi T, Jalalian S, Weickenmeier J (2021) Brain aging mechanisms with mechanical manifestations. Mech Ageing Dev 200:111575.

Burt JB, Helmer M, Shinn M, Anticevic A, Murray JD (2020) Generative modeling of brain maps with spatial autocorrelation. NeuroImage 220:117038.

Casey B, Tottenham N, Liston C, Durston S (2005) Imaging the developing brain: what have we learned about cognitive development? Trends Cogn Sci 9:104–110.

Chen G, Saad ZS, Britton JC, Pine DS, Cox RW (2013) Linear mixed-effects modeling approach to FMRI group analysis. NeuroImage 73:176–190.

Chettouf S, Triebkorn P, Daffertshofer A, Ritter P (2022) Unimanual sensorimotor learning—A simultaneous EEG-FMRI aging study. Hum Brain Mapp 43:2348–2364.

Cleveland WS (1981) LOWESS: a program for smoothing scatterplots by robust locally weighted regression. Am Stat 35:54.

Cordes D, Haughton VM, Arfanakis K, Carew JD, Turski PA, Moritz CH, Quigley MA, Meyerand ME (2001) Frequencies contributing to functional connectivity in the cerebral cortex in “resting-state” data. Am J Neuroradiol 22:1326–1333.

Coupé P, Catheline G, Lanuza E, Manjón JV, Alzheimer’s Disease Neuroimaging Initiative (2017) Towards a unified analysis of brain maturation and aging across the entire lifespan: A MRI analysis. Hum Brain Mapp 38:5501–5518.

Deery HA, Di Paolo R, Moran C, Egan GF, Jamadar SD (2023) The older adult brain is less modular, more integrated, and less efficient at rest: A systematic review of large-scale resting-state functional brain networks in aging. Psychophysiol 60:e14159.

Demirtaş M, Burt JB, Helmer M, Ji JL, Adkinson BD, Glasser MF, Van Essen DC, Sotiropoulos SN, Anticevic A, Murray JD (2019) Hierarchical heterogeneity across human cortex shapes large-scale neural dynamics. Neuron 101:1181–1194.

Dumontheil I, Burgess PW, Blakemore S (2008) Development of rostral prefrontal cortex and cognitive and behavioural disorders. Dev Med Child Neurol 50:168–181.

Edde M, Leroux G, Altena E, Chanraud S (2021) Functional brain connectivity changes across the human life span: from fetal development to old age. J Neurosci Res 99:236–262.

Fabiani M, Gordon BA, Maclin EL, Pearson MA, Brumback-Peltz CR, Low KA, McAuley E, Sutton BP, Kramer AF, Gratton G (2014) Neurovascular coupling in normal aging: A combined optical, ERP and fMRI study. NeuroImage 85:592–607.

Faghiri A, Stephen JM, Wang Y-P, Wilson TW, Calhoun VD (2019) Brain development includes linear and multiple nonlinear trajectories: a cross-sectional resting-state functional magnetic resonance imaging study. Brain Connect 9:777–788.

Fair DA, Cohen AL, Power JD, Dosenbach NUF, Church JA, Miezin FM, Schlaggar BL, Petersen SE (2009) Functional brain networks develop from a “local to distributed” organization. PLoS Comput Biol 5:e1000381.

Fallon J, Ward PGD, Parkes L, Oldham S, Arnatkevičiūtė A, Fornito A, Fulcher BD (2020) Timescales of spontaneous fMRI fluctuations relate to structural connectivity in the brain. Netw Neurosci 4:788–806.

Fascianelli V, Tsujimoto S, Marcos E, Genovesio A (2019) Autocorrelation structure in the macaque dorsolateral, but not orbital or polar, prefrontal cortex predicts response-coding strength in a visually cued strategy task. Cereb Cortex 29: 230–241.

Ferreira LK, Busatto GF (2013) Resting-state functional connectivity in normal brain aging. Neurosci Biobehav Rev 37:384–400.

Fischl B (2012) FreeSurfer. NeuroImage 62:774–781.

Flechsig P (1901) Developmental (myelogenetic) localisation of the cerebral cortex in the human subject. Lancet 158:1027–1030.

Fransson P, Åden U, Blennow M, Lagercrantz H (2011) The functional architecture of the infant brain as revealed by resting-state fMRI. Cereb Cortex 21:145–154.

Friston KJ, Holmes AP, Poline J-B, Grasby PJ, Williams SCR, Frackowiak RSJ, Turner R (1995) Analysis of fMRI time-series revisited. NeuroImage 2:45–53.

Ganzetti M, Wenderoth N, Mantini D (2014) Whole brain myelin mapping using T1- and T2-weighted MR imaging data. Front Hum Neurosci 8:671.

Gao R, Van Den Brink RL, Pfeffer T, Voytek B (2020) Neuronal timescales are functionally dynamic and shaped by cortical microarchitecture. eLife 9:e61277.

Geerligs L, Tsvetanov KA, Cam-CAN, Henson RN (2017) Challenges in measuring individual differences in functional connectivity using fMRI: The case of healthy aging. Hum Brain Mapp 38:4125–4156.

Glasser MF, Smith SM, Marcus DS, Andersson JLR, Auerbach EJ, Behrens TEJ, Coalson TS, Harms MP, Jenkinson M, Moeller S, Robinson EC, Sotiropoulos SN, Xu J, Yacoub E, Ugurbil K, Van Essen DC (2016) The Human Connectome Project’s neuroimaging approach. Nat Neurosci 19:1175–1187.

Glasser MF, Sotiropoulos SN, Wilson JA, Coalson TS, Fischl B, Andersson JL, Xu J, Jbabdi S, Webster M, Polimeni JR, Van Essen DC, Jenkinson M (2013) The minimal preprocessing pipelines for the Human Connectome Project. NeuroImage 80:105–124.

Glasser MF, Van Essen DC (2011) Mapping human cortical areas in vivo based on myelin content as revealed by T1- and T2-Weighted MRI. J Neurosci 31:11597–11616.

Gogtay N, Giedd JN, Lusk L, Hayashi KM, Greenstein D, Vaituzis AC, Nugent TF, Herman DH, Clasen LS, Toga AW, Rapoport JL, Thompson PM (2004) Dynamic mapping of human cortical development during childhood through early adulthood. Proc Natl Acad Sci USA 101:8174–8179.

Golshani P, Gonçalves JT, Khoshkhoo S, Mostany R, Smirnakis S, Portera-Cailliau C (2009) Internally mediated developmental desynchronization of neocortical network activity. J Neurosci 29:10890–10899.

Gribizis A, Ge X, Daigle TL, Ackman JB, Zeng H, Lee D, Crair MC (2019) Visual cortex gains independence from peripheral drive before eye opening. Neuron 104:711–723.

Grydeland H, Vértes PE, Váša F, Romero-Garcia R, Whitaker K, Alexander-Bloch AF, Bjørnerud A, Patel AX, Sederevičius D, Tamnes CK, Westlye LT, White SR, Walhovd KB, Fjell AM, Bullmore ET (2019) Waves of maturation and senescence in micro-structural MRI markers of human cortical myelination over the lifespan. Cereb Cortex 29:1369–1381.

Han S, Zheng R, Li S, Zhou B, Jiang Y, Wang C, Wei Y, Pang J, Li H, Zhang Y, Chen Y, Cheng J (2022) Integrative functional, molecular, and transcriptomic analyses of altered intrinsic timescale gradient in depression. Front Neurosci 16:826609.

Harada CN, Natelson Love MC, Triebel KL (2013) Normal cognitive aging. Clin Geriatr Med 29:737–752.

Harms MP et al. (2018) Extending the human connectome project across ages: imaging protocols for the lifespan development and aging projects. NeuroImage 183:972–984.

Honey CJ, Thesen T, Donner TH, Silbert LJ, Carlson CE, Devinsky O, Doyle WK, Rubin N, Heeger DJ, Hasson U (2012) Slow cortical dynamics and the accumulation of information over long timescales. Neuron 76:423–434.

Huang Z, Liu X, Mashour GA, Hudetz AG (2018) Timescales of intrinsic BOLD signal dynamics and functional connectivity in pharmacologic and neuropathologic states of unconsciousness. J Neurosci 38:2304–2317.

Huntenburg JM, Bazin P-L, Margulies DS (2018) Large-scale gradients in human cortical organization. Trends Cogn Sci 22:21–31.

Ito T, Hearne LJ, Cole MW (2020) A cortical hierarchy of localized and distributed processes revealed via dissociation of task activations, connectivity changes, and intrinsic timescales. NeuroImage 221:117141.

Khazipov R, Luhmann HJ (2006) Early patterns of electrical activity in the developing cerebral cortex of humans and rodents. Trends Neurosci 29:414–418.

Kiebel SJ, Daunizeau J, Friston KJ (2008) A Hierarchy of time-scales and the brain. PLoS Comput Biol 4: e1000209.

Levy R (1994) Aging-associated cognitive decline. Int Psychogeriatr 6:63–68.

Li J, Kong R, Liégeois R, Orban C, Tan Y, Sun N, Holmes AJ, Sabuncu MR, Ge T, Yeo BTT (2019) Global signal regression strengthens association between resting-state functional connectivity and behavior. NeuroImage 196:126–141.

Lippe S, Roy M-S, Perchet C, Lassonde M (2006) Electrophysiological markers of visuocortical development. Cereb Cortex 17:100–107.

Manea AM, Zilverstand A, Ugurbil K, Heilbronner SR, Zimmermann J (2022) Intrinsic timescales as an organizational principle of neural processing across the whole rhesus macaque brain. eLife 11:e75540.

Manea AMG, Maisson DJ-N, Voloh B, Zilverstand A, Hayden B, Zimmermann J (2024) Neural timescales reflect behavioral demands in freely moving rhesus macaques. Nat Commun 15:2151.

Martini FJ, Guillamón-Vivancos T, Moreno-Juan V, Valdeolmillos M, López-Bendito G (2021) Spontaneous activity in developing thalamic and cortical sensory networks. Neuron 109:2519–2534.

Murman D (2015) The impact of age on cognition. Semin Hear 36:111–121.

Murphy K, Birn RM, Bandettini PA (2013) Resting-state fMRI confounds and cleanup. NeuroImage 80:349–359.

Murphy K, Fox MD (2017) Towards a consensus regarding global signal regression for resting state functional connectivity MRI. NeuroImage 154:169–173.

Murray JD, Bernacchia A, Freedman DJ, Romo R, Wallis JD, Cai X, Padoa-Schioppa C, Pasternak T, Seo H, Lee D, Wang X-J (2014) A hierarchy of intrinsic timescales across primate cortex. Nat Neurosci 17:1661–1663.

Power JD, Barnes KA, Snyder AZ, Schlaggar BL, Petersen SE (2012) Spurious but systematic correlations in functional connectivity MRI networks arise from subject motion. NeuroImage 59:2142–2154.

Raut RV, Snyder AZ, Raichle ME (2020) Hierarchical dynamics as a macroscopic organizing principle of the human brain. Proc Natl Acad Sci USA 117:20890–20897.

Salthouse TA (2010) Selective review of cognitive aging. J Int Neuropsychol Soc 16:754–760.

Schaefer A, Kong R, Gordon EM, Laumann TO, Zuo X-N, Holmes AJ, Eickhoff SB, Yeo BTT (2018) Local-global parcellation of the human cerebral cortex from intrinsic functional connectivity MRI. Cereb Cortex 28:3095–3114.

Schmithorst VJ, Vannest J, Lee G, Hernandez-Garcia L, Plante E, Rajagopal A, Holland SK, The CMIND Authorship Consortium (2015) Evidence that neurovascular coupling underlying the BOLD effect increases with age during childhood: neuronal-vascular coupling increases with age. Hum Brain Mapp 36:1–15.

Sela RJ, Hurvich CM (2009) Computationally efficient methods for two multivariate fractionally integrated models. J Time Ser Anal 30: 631–651.

Sethi SS, Zerbi V, Wenderoth N, Fornito A, Fulcher BD (2017) Structural connectome topology relates to regional BOLD signal dynamics in the mouse brain. Chaos 27:047405.

Shafiei G, Markello RD, Vos De Wael R, Bernhardt BC, Fulcher BD, Misic B (2020) Topographic gradients of intrinsic dynamics across neocortex. eLife 9:e62116.

Shinn M, Hu A, Turner L, Noble S, Preller KH, Ji JL, Moujaes F, Achard S, Scheinost D, Constable RT, Krystal JH, Vollenweider FX, Lee D, Anticevic A, Bullmore ET, Murray JD (2023) Functional brain networks reflect spatial and temporal autocorrelation. Nat Neurosci 26:867–878.

Siegle JH et al. (2021) Survey of spiking in the mouse visual system reveals functional hierarchy. Nature 592:86–92.

Song HF, Kennedy H, Wang X-J (2014) Spatial embedding of structural similarity in the cerebral cortex. Proc Natl Acad Sci USA 111:16580–16585.

Song M, Shin EJ, Seo H, Soltani A, Steinmetz NA, Lee D, Jung MW, Paik S-B (2024) Hierarchical gradients of multiple timescales in the mammalian forebrain. Proc Natl Acad Sci USA 121:e2415695121.

Sowell ER, Thompson PM, Leonard CM, Welcome SE, Kan E, Toga AW (2004) Longitudinal mapping of cortical thickness and brain growth in normal children. J Neurosci 24:8223–8231.

Springer SD, Erker TD, Schantell M, Johnson HJ, Willett MP, Okelberry HJ, Rempe MP, Wilson TW (2023) Disturbances in primary visual processing as a function of healthy aging. NeuroImage 271:120020.

Stiles J, Jernigan TL (2010) The basics of brain development. Neuropsychol Rev 20:327–348.

Supekar K, Uddin LQ, Prater K, Amin H, Greicius MD, Menon V (2010) Development of functional and structural connectivity within the default mode network in young children. NeuroImage 52:290–301.

Sydnor VJ, Larsen B, Bassett DS, Alexander-Bloch A, Fair DA, Liston C, Mackey AP, Milham MP, Pines A, Roalf DR, Seidlitz J, Xu T, Raznahan A, Satterthwaite TD (2021) Neurodevelopment of the association cortices: Patterns, mechanisms, and implications for psychopathology. Neuron 109:2820–2846.

Sydnor VJ, Larsen B, Seidlitz J, Adebimpe A, Alexander-Bloch AF, Bassett DS, Bertolero MA, Cieslak M, Covitz S, Fan Y, Gur RE, Gur RC, Mackey AP, Moore TM, Roalf DR, Shinohara RT, Satterthwaite TD (2023) Intrinsic activity development unfolds along a sensorimotor–association cortical axis in youth. Nat Neurosci 26:638–649.

Takesian AE, Hensch TK (2013) Balancing plasticity/stability across brain development. Prog Brain Res 207:3–34.

Thomas Yeo BT, Krienen FM, Sepulcre J, Sabuncu MR, Lashkari D, Hollinshead M, Roffman JL, Smoller JW, Zöllei L, Polimeni JR, Fischl B, Liu H, Buckner RL (2011) The organization of the human cerebral cortex estimated by intrinsic functional connectivity. J Neurophysiol 106:1125–1165.

Truzzi A, Cusack R (2023) The development of intrinsic timescales: A comparison between the neonate and adult brain. NeuroImage 275:120155.

Tsvetanov KA, Henson RNA, Rowe JB (2021) Separating vascular and neuronal effects of age on fMRI BOLD signals. Phil Trans R Soc Lond B Biol Sci 376:20190631.

Uscătescu LC, Kronbichler M, Said-Yürekli S, Kronbichler L, Calhoun V, Corbera S, Bell M, Pelphrey K, Pearlson G, Assaf M (2023) Schizophrenia 9:18.

Watanabe T, Rees G, Masuda N (2019) Atypical intrinsic neural timescale in autism. eLife 8:e42256.

West KL, Zuppichini MD, Turner MP, Sivakolundu DK, Zhao Y, Abdelkarim D, Spence JS, Rypma B (2019) BOLD hemodynamic response function changes significantly with healthy aging. NeuroImage 188:198–207.

Wolff A, Berberian N, Golesorkhi M, Gomez-Pilar J, Zilio F, Northoff G (2022) Intrinsic neural timescales: temporal integration and segregation. Trends Cogn Sci 26:159–173.

Wu K and Gollo LL (2025) Mapping and modeling age-related changes in intrinsic neural timescales Commun Biol 8:167.

Zhang A, Wengler K, Zhu X, Horga G, Goldberg TE, Lee S, and For Alzheimer’s Disease Neuroimaging Initiative. Altered hierarchical gradients of intrinsic neural timescales in mild cognitive impairment and Alzheimer’s disease. J Neurosci 44: e2024232024.

